# Rapid Vascular Activation Precedes Immune Cell Infiltration Following Corneal Alkali Burn

**DOI:** 10.64898/2026.08.21.746379

**Authors:** Naoufal Akla, Marc Groleau, Meagan Jade Latorre, Grace Lin, Delali Shana Degue, Marie-Claude Robert, Bruno Larrivée, Christopher E. Rudd, May Griffith

## Abstract

Under homeostatic conditions, the cornea is avascular and contains few immune cells, but this changes rapidly following injury. Although the long-term consequences of corneal damage are well characterized, the earliest vascular and immune responses remain poorly understood. Here, we used a murine corneal alkali-burn model to examine limbal vascular activation and leukocyte recruitment immediately and at 2, 6, and 24 hours after injury. Limbal blood vessels underwent immediate dilation; however, vascular leakage into the corneal stroma occurred only in males. Lymphatic capillaries rapidly formed directed extensions toward the injury without significantly increasing their total vascular area, with males exhibiting longer extensions than females. Fluorescent dextran uptake provided evidence that these lymphatic vessels were functionally engaged in early tracer drainage. Despite pronounced vascular activation, early recruitment of neutrophils, monocytes, dendritic cells, macrophages, T cells, B cells, and natural killer cells remained limited. Thus, limbal blood and lymphatic vessels initiate the earliest response to corneal alkali injury before substantial leukocyte infiltration. These findings reveal sex-dependent differences in vascular permeability and lymphatic remodeling and identify the limbal vasculature as an early regulator of corneal inflammation and tissue repair.

## I. Introduction

The cornea is the transparent front of the eye and main refracting lens focusing incoming light for vision. Its external location renders it susceptible to accidental injuries, including chemical burns. Chemical burns cause 11.5% to 22.1% of eye injuries and 60% of those are alkali burns (1). These burns often result from accidental exposure at work or to household cleaning products (2).Acid burns typically cause superficial coagulation, but alkali burns rapidly penetrate deep into tissues due to their lipophilicity (3). The hydroxyl anions (OH,) of alkalis saponify fatty acids permitting the chemical to cross cell membranes, followed by precipitation and denaturation of proteoglycans and stromal collagen allowing the alkali to penetrate further in the eye (4, 5).

Common alkalis implicated in eye injuries include ammonia (NH^3^), potassium hydroxide (KOH) and sodium hydroxide (NaOH) (1). In ophthalmology, controlled alkali burns using NaOH are routinely used to model severe corneal pathologies as they trigger immediate inflammation and even after healing, render the ocular surface susceptible to auto-immune-like conditions (6). Alkali exposure results in acute inflammation and a vascular response in which the blood flow to the inflamed area increases due vasodilation at the injury site. Capillary permeability increases due to endothelial barrier disruption, mainly from junctional gaps and transcytosis mechanisms that can result in edema (7). This will be accompanied by cellular events where leukocytes, such as neutrophils, will be recruited into the injured area.

In this context, the immune system is comprised of multiple specialized cell types that work together to protect the host against viral infections and contribute to tissue repair and wound healing. In T-cells, a complex network of surface receptors and iintracellular signaling pathways are triggered upon antigen receptor (TCR) engagement (8–11). Central to the activation process are the the CD4 or CD8 co-receptors which associate with the Src-family kinase p56^lck^ (11, 12) and phosphorylates immunoreceptor tyrosine-based activation motifs (ITAMs) in TCRζ/CD3 subunits initiating downstream signaling (13–16). This involves the recruitment of a second tyrosine kinase, termed ζ-chain-associated protein kinase 70 (ZAP-70) (17) and the phosphorylation of numerous downstream enzymes and adaptor proteins (18–23). The ability to T-cells to migrate to sites of infection or injury depend in these activation events. Other immune cells such as B and mast cells also use related src kinases to initiate their activation.

Following tissue injury, innate and adaptive immune cells coordinate inflammatory responses in a process involving the recruitment of various leukocyte populations (24–26). Neutrophils are generally seen to be the earliest cells recruited to damaged tissues, where they contribute to microbial defence and can influence vascular permeability. Infiltrating monocytes have been reported to differentiate into macrophages or dendritic cells (DCs), and as such, are involved in debris clearance and antigen presentation (24, 26). In particular, DCs coordinate innate and adaptive responses by transporting antigens through lymphatic vessels to present antigen to local lymph nodes. In this regard, macrophages can either amplify inflammation, or facilitate resolution depending on their activation state (27, 28). T and B cells as well as natural killer (NK) cells can provide additional cytotoxic functions, and in this sense, can modulate the magnitude of corneal inflammation (29–31).

Although cellular responses have been studied during established corneal inflammation, less is known about the early kinetics following alkali injury and whether leukocyte recruitment occurs simultaneously with the activation of limbal blood and lymphatic vessels. We therefore sought to define the immediate vascular and cellular events occurring during the first 24 hours following NaOH-induced corneal alkali burn. Using morphological, functional and flow cytometric approaches, we have examined the temporal responses of limbal blood vessels, lymphatic vessels and the infiltrating immune-cell populations. Our findings show that corneal alkali injury triggers an immediate phase dominated by rapid vasodilation, increased vascular permeability and functional lymphatic activation, events which precede substantial leukocyte recruitment. These observations identify the limbal vasculature as an active initiator of the inflammatory response and redefine the earliest phase of corneal alkali injury as one characterized primarily by vascular rather than immune cell infiltration.

## II. Methods

### Mouse corneal alkali burns

CD1 outbred female and male mice, 8 weeks or older (Charles River, Montreal, CA), were used in accordance with the ARRIVE guidelines for reporting animal research. Animal protocols were approved by “Le Comité de protection des animaux du CIUSSS de l’Est-de-l’Île-de-Montréal (CPA-CEMTL)” (Protocol 2021-2356 and 2025-3858). Each mouse was given a general anaesthetic of isoflurane. An alkali burn was created in the centre of each right cornea (OD) by placing a 2 mm diameter filter paper disc saturated with 0, 0.125, 0.250, and 0.5N NaOH (Aldrich, Oakville, Canada) for 15 seconds, after which the eye was flushed with phosphate buffered saline (PBS) to remove any remaining chemical. The contralateral untreated left corneas (OS) served as controls (Fig. 1a). Tetracaine hydrochloride (0.5%, Minims, Bausch and Lomb, Vaughn, Canada) and buprenorphine (0.05 mg/Kg) (Vetergesic, CDMV Inc., Saint-Hyacinthe, Canada) were used for local and systemic analgesia.

**Fig. 1.**
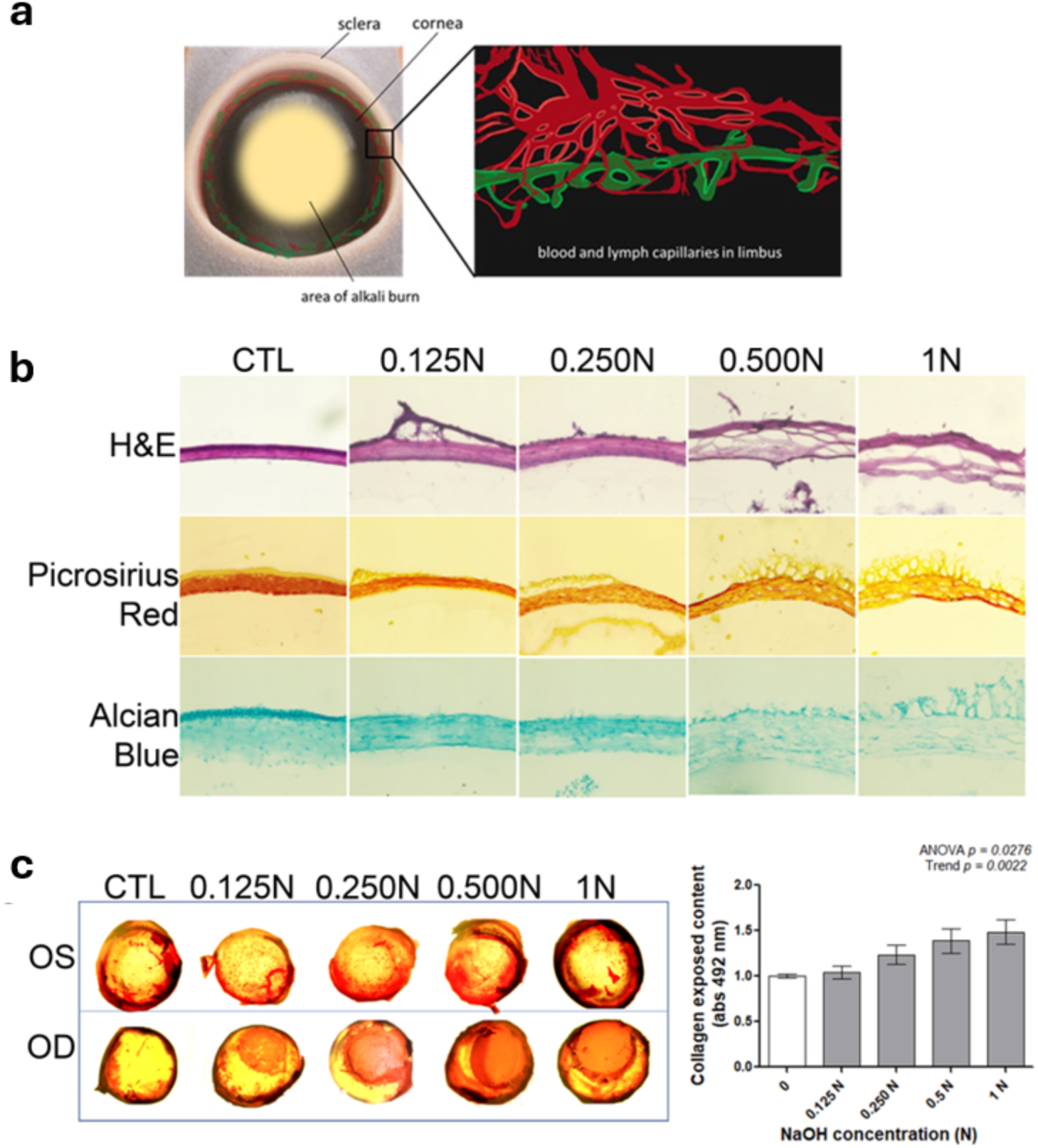
Alkali burned mouse corneas. a) Location of alkali burn in the central cornea and high magnification of the limbal region showing the lymph and blood capillaries in a healthy, normal cornea. b) Representative sections of mouse corneas exposed to increasing concentrations of NaOH (0.125N to 1N) stained with H&E (top), Picrosirius Red (middle) and Alcian Blue (bottom). c) Wholemounts of representative Picrosirius Red stained burned right (OD) corneas and the contralateral untreated left (OS) corneas (n=3 per group) showing exposed collagen that is quantified by dye elution and absorbance at 492 nm (right panel). One-way ANOVA followed by a post hoc linear trend was used to assess dose–response significance (ANOVA p = 0.028; trend p = 0.002). Bar graph represents mean ± SEM; of at least n = 3 corneas/group. p < 0.05 compared to control; NS: not significant.

Postoperative observation of the anterior segment was performed immediately (T=0 hour), 2, 6 and 24 hours post corneal burn on mouse eyes. The thickness of the central cornea and presence of edema were assessed using optical coherence tomography (OCT) (Lumedica, OQ LabScope). Epithelial integrity was evaluated by corneal fluorescein staining (CFS), where intact epithelium excludes the dye but defects allow dye entry, with fluorescence captured on the MICRON IV system (Phoenix Research Labs, Pleasanton, CA, USA). The untreated contralateral eyes served as controls. After each examination, mice were euthanized and whole treated and untreated eyes were fixed in and processed based on the methodology below.

### Histopathology

Excised corneas with scleral rims were embedded in OCT (Thermo Fisher, Waltham, MA) and frozen for cryosectioning. Representative sections were stained with Haematoxylin and Eosin (H&E) for histopathological examination. To evaluate the presence of collagen fibers, picrosirius red staining was performed using a picrosirius stain kit (ab150681, Abcam, Cambridge, UK). To determine proteoglycans content, samples were incubated in Alcian blue solution for 30 minutes.

### Exposed corneal collagen content by Picrosirius Red staining

After alkali burn was induced, exposed corneal collagen was characterized by Picrosirius Red staining (1 mg/mL Sirius Red saturated Picric acid) (32, 33). Briefly, post overnight fixation in PFA 1% of whole eye globe, excised corneas were incubated for 1.5 hours at room temperature (RT). The stained samples were thoroughly washed with distilled water to remove unbound dye and dried at 37°C before qualitative analysis by stereomicroscopy EVOS-XL core. For quantitative analysis, the Picrosirius Red dye was dissolved in 0.2M NaOH/methanol (1:1 ratio) in 250 µl with mild shaking 300 rpm for 1h at RT, and the optical density was measured at 492 nm using a microplate reader (Tecan Spark, Thermo Fisher Scientific, Ste-Laurent, Canada) for duplicate samples for each condition.

### Immunohistochemistry

Immunohistochemistry was performed according to Anderson (2014) with modifications (34). Briefly, freshly excised eyeballs were fixed overnight in buffered 2% PFA at 4°C and corneas were excised for immunofluorescence staining and mounted. Nonspecific staining was blocked in 5% goat serum 0.5% Triton X,100 in PBS (BD Biosciences, San Jose, CA). Samples were stained with rabbit polyclonal anti–mouse CD31 antibody (Ab28364) for blood vessels alone, or double stained with rat anti–mouse CD31 monoclonal antibody (BD553370) and rabbit anti–mouse LYVE-1 antibody (Ab218535) for blood and lymphatic vessels. Primary antibodies were visualized by fluorescent tagged matching goat anti–rat (A11007) or goat anti-rabbit secondary antibodies (A11034). Stained samples were cut in 4 equal quadrant and covered with Fluoroshield mounting medium (F6057) then examined by epifluorescence microscopy (EVOS FL or XL Thermofisher). Vascular structures stained as CD31+LYVE-1− were identified as blood vessels, whereas those stained as CD31+LYVE-1+ were defined as lymphatic vessels.

### Flatmounts for vascular assessments

Four radial cuts were made to flatten the corneas, which were then stained with anti-CD31 and anti-LYVE-1 as described above to label blood and lymphatic vessels respectively. The labelled vessels were then analyzed using the ImageJ software (National Institutes of Health available at http://rsb.info.nih.gov/ij/index.html). To calculate the average microvessel density, the semi-automated Vesselj plug-in to quantify the ratio of vascularized area to total corneal-limbal plexus was used (35). Conjunctival and ciliary vessels that terminate and form loops in the limbus are often termed arcades and located near the corneal border. A custom ImageJ/Fiji macro called VasoMetrics was used to measure blood vessel diameters as orthogonal distances from centerlines drawn across the innermost final end-loops of the limbal marginal capillary arcade bordering the peripheral cornea. Measurements were set at 10 um intervals (36). Limbal lymphatic vessels area coverage around the corneal edge were measured from centered images around the most proximal corneo-limbal lymphatic arcade including their extension toward the corneal center excluding afferent ducts (37). The length of that extension was measured by drawing a line from the blind-ended lymphatic capillary tip perpendicularly to the most proximal originating corneo-limbal arcade.

### Lymphatic vessel function

To determine whether the extended lymphatic vessels entering the cornea immediately after a burn were functional or not, a 40 kDa dextran-FITC dye was applied into the 0.25N NaOH wound. The passage of the fluorescent signal from initial uptake into lymphatic capillaries and subsequent clearance through collecting vessels and cervical lymph nodes (CLN’s) was tracked using Micron IV imaging using the fluorescence mode (38). The dye molecules are too large to enter blood capillaries and can only be cleared by lymphatic vessels. Fluorescent FITC-Dextran interstitial tracer initial lymphatic uptake was assessed with modifications as described in (27, 28). Briefly, 2 uL of (FITC) Dextran (40 kDa) at a concentration of 2 mg/ml was topically applied to the injured eye surface 1 to 30 minutes before sacrifice (contralateral eye served as unwounded control). Images were taken at 5, 10, and 20 minutes post inoculation before sacrifice. Blood was extracted and excised CLN’s immediately fixed in 4% PFA and imbedded in optimal cutting temperature O.C.T cryopreserved for cryosections (39). Clearance was assessed by quantifying the surface area percentage of CLN sections fluorescence acquired at 1- and 30-minutes intervals (40).

### Vascular permeability Assay

Permeability of the vessels were assessed by Evans Blue perfusion as previously described (38, 41). Evans blue was injected at 45 mg/Kg into the mice by tail vein and circulated for 2 hours prior to sacrifice and corneal excision. The excised corneas were incubated with N,N,dimethylformamide for 24 h at 65 °C then centrifuged for 45 minutes at 13300 rpm. Evans Blue-Albumin complex (EBA) was quantified shortly after by fluorometry at 620 nm maximal absorbance and corrected with 740 nm minimum absorbance background using TECAN Spark. Through standard curve established by known EBA concentrations corneal tissue content were measured in ug/mL/cornea.

### Flow Cytometric Analysis of Corneal Immune Cells

CD1 mice were given alkali burns, with three concentrations of NaOH, as explained earlier. For a negative control, some mice were treated with 0.000N NaOH. After 0, 2, 6, and 24 hours, both eyes were enucleated and immediately dissected to isolate the cornea. The sclera, conjunctiva, and limbus were all removed when isolating the cornea. Corneas were then incubated in 1 mg/mL of collagenase from Clostridium histolyticum, type XI (C9407-100MG, Sigma-Aldrich, Oakville, Canada) for 2 hours at 37°C with constant agitation. Following the collagen digestion, cells were passed through a 70µm cell strainer to obtain a single cell suspension. Cells were stained for 20 minutes with a fixable viability dye (564406, Becton, Dickinson and Company, Mississauga, Canada). Cells were then treated with Fc block (501129393, FisherSci, Saint-Laurent, Canada) for 30 minutes to reduce non-specific antibody staining. Finally, cells were stained with surface antibodies to identify various immune cells (Supplemental Table S1). Samples were then fixed in 4% PFA in PBS for 20 minutes and resuspended in a solution containing Precision Count Beads™ (424902, BioLegend, San Diego, USA). Samples were then acquired on a LSRFortessa™ X-20 (Becton, Dickinson and Company, Mississauga, Canada). Data was analyzed on FlowJo (v10.10.1, Becton, Dickinson and Company, Mississauga, Canada). Immune cell populations were made using a combination of fluorescence minus one (FMO) as negative controls and other tissues, including spleen, lungs, bone marrow, and a peritoneal lavage, as positive controls. Total immune cells were identified by being CD45+ and were further categorized as: Neutrophils (Ly-6C+, Ly-6G+), monocytes (Ly-6C+, Ly-6G-, CD11b+), dendritic cells (CD11c+, CD11b+), macrophages (F4/80+, CD11b+), T cells (CD3+), B cells (CD19+), and NK cells (NK1.1+). Flow cytometric gating strategy for immune cell populations are shown in Supplemental figures 1 and 2.

### Statistical Analysis

Data are expressed as means ± standard error of the mean (SEM). Statistical analyses were performed using GraphPad Prism software (version 5.0, GraphPad Software, San Diego, CA, USA). CTL groups were made by pooling the results of non-injured contralateral corneas together. Comparisons between two groups were conducted using an unpaired Student’s t-test. For comparisons involving more than two groups, one-way analysis of variance (ANOVA) followed by Tukey’s or Trend post hoc test was used to identify specific group differences. Statistical significance was set at a p-value < 0.05. Sample sizes and the specific statistical tests employed are detailed in each corresponding figure legend.

## III. Results

### Gross changes after alkali exposure

Exposure to NaOH results in an area of opacity in the central cornea (**Fig. 1a**), away from the limbal area where blood vessels and lymphatics are located. With increasing concentrations, swelling of the cornea results from disruption of the corneal epithelium and stroma (**Fig. 1b**). Picrosirius Red staining showed increased collagen fiber disorganization and exposure with increased NaOH concentration. Conversely, proteoglycans loss confirmed matrix damage and collagen fiber exposure (**Fig. 1c**). Burns caused by 0.125N and 0.25N NaOH showed immediate epithelial damage as revealed by fluorescein uptake but recovered within 24 hours (**Fig. 2a-c**). However, corneas exposed to 0.50N did not heal over the 24 hours (**Fig. 2a, d**). OCT observations showed at the lowest concentration of 0.125N, corneal swelling was seen at two hours after NaOH exposure but returned to a thickness comparable to the control by 24 hours (**Fig. 2e**). For higher concentrations, swelling was immediate and remained over the 24-hour observation period.

**Fig. 2.**
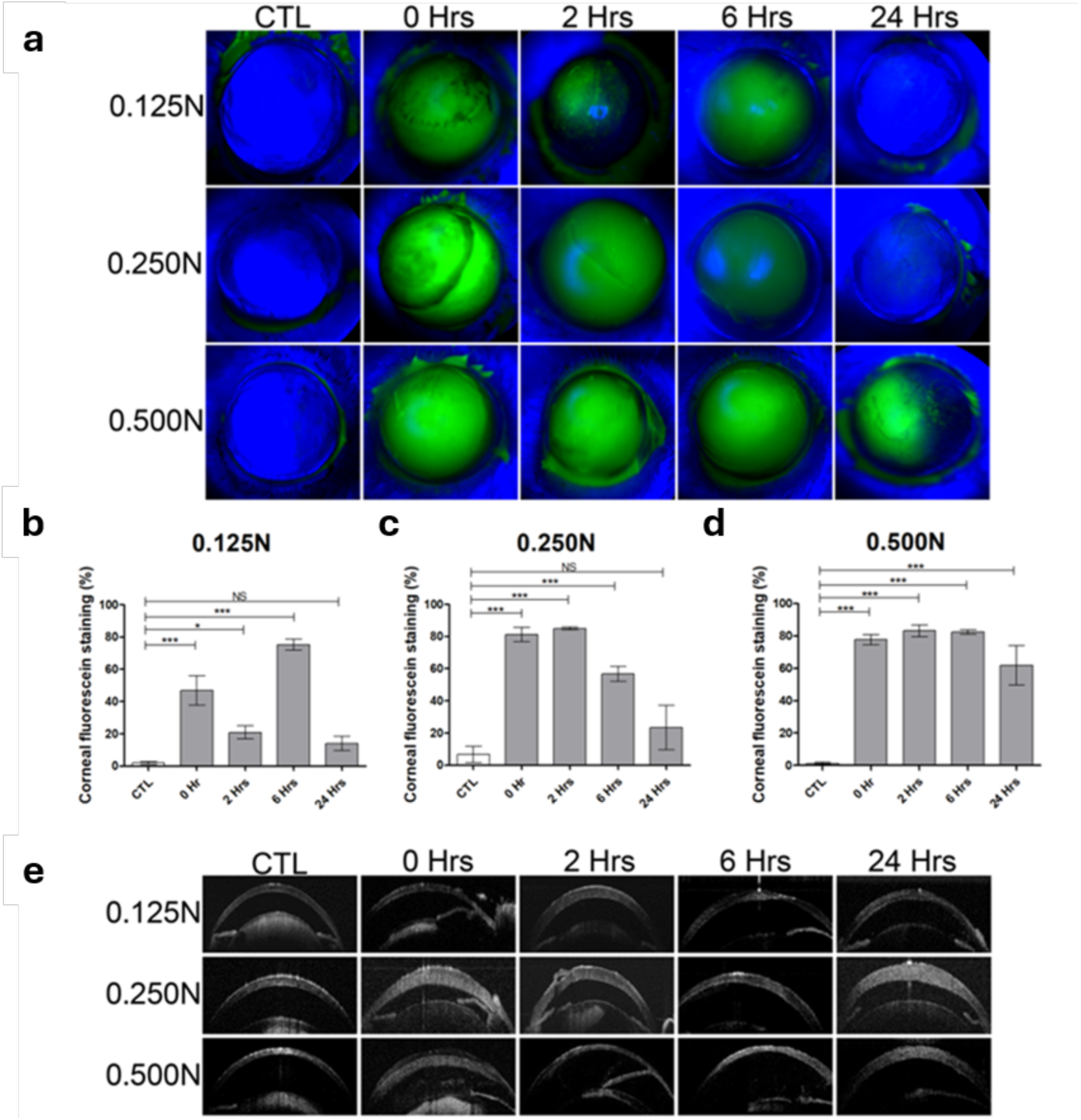
Characteristics of the *in vivo* cornea after alkali burn injury. a) Representative fluorescein-stained cornea images at 0, 2, 6 and 24 hours post-injury. b-d) Quantification of corneal fluorescein staining percentage of area fractions of the burned area as analyzed with ImageJ. e) Representative OCT images of the cornea and anterior segment showing changes in corneal thickness over time after exposure to 0.125N, 0.25N and 0.5N NaOH. One-way ANOVA followed by Tukey’s test was used to compare each time point to pooled controls (OS). Asterisks indicate significant post hoc differences: *p < 0.05; **p < 0.01; ***p < 0.001; ****p < 0.0001; NS = not significant.

### Changes to limbal blood capillaries

Examination of red CD31-stained marginal limbal arcade loop capillaries showed that exposure to 0.125N NaOH did not result in any observable morphological changes (**Fig. 3a**). Exposure to 0.25N and 0.5N showed more red fluorescence. At these higher doses, morphometric analyses showed that the area occupied by limbal blood vessels was significantly reduced at specific later time points, whereas the lowest dose had no effect (**Fig. 3b-d)**. Further analyses of capillary loop diameters showed an overall progressive increase in diameters overtime after the burn throughout all the doses tested, which were significant immediately after injury (T=0 hour) (**Fig. 3e**). Surprisingly, at 0.125N, no immediate latent phase was observed, the response appeared biphasic whereas 0.25N exposure showed monophasic over time. As expected, the higher concentration of 0.5N exhibited a longer latent phase, a phenomenon observed before the onset of angiogenesis as it transitions into the proliferative stage during the aberrant wound healing response. The most consistent increase occurred with 0.25N, showing a significant and sustained amplitude relative to controls throughout the 24 hours kinetic. At that endpoint, all doses demonstrated a significant increase in capillaries diameters, nearly doubling control values.

**Fig. 3.**
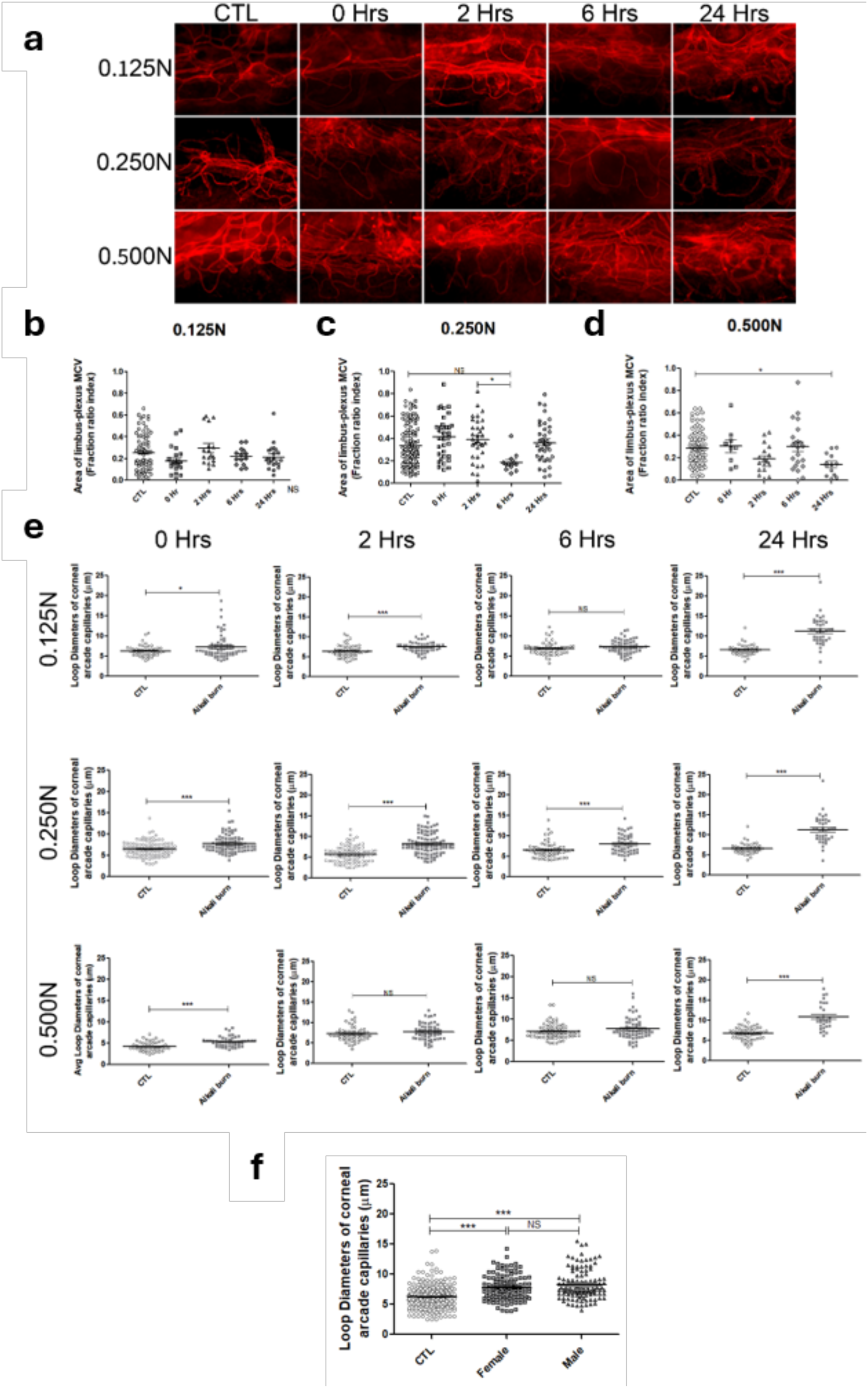
Early blood vascular events in alkali induced acute inflammatory response. a) Representative 20X flatmount CD31 staining images of limbal microvasculature after NaOH injury (0.125N, 0.25N, 0.5N) at indicated time points. b,c,d) Quantification of limbal microvascular network area using VesselJ plugin and ImageJ thresholding of an average of 5 images per cornea across 4 quadrants. Data are presented as mean ± SEM; with individual points representing each image. n = 3 mice for most of the analysis, except for the 0.125N/CTL group (n = 11), 0.25N/24h group (n = 4), 0.25N/CTL group (n = 13), 0.5N/0h group (n = 2), and 0.5N/CTL group (n = 11). One Way ANOVA with Tukey post-hoc test was used for vessel Area. ANOVA b) p = 0.0960; c) p = 0.0029; d) p < 0.0096. Asterisks indicate significant post hoc differences: *p < 0.05; **p < 0.01; ***p < 0.001; NS = not significant. e) Quantification of loop diameters from the most marginal arcade capillaries at the corneal periphery using the VasoMetrics ImageJ macro, measuring distances orthogonal to vessel centerlines. allowing all values across the entire vessels to be averaged together for a single, representative diameter of a given loop segment. An average of 3 loops measurements across 8 fields of view per cornea. Student T test was used to compare groups. Each point represents the average of each loop presented as an overall mean ± SEM; n = 3 mice for most of the analysis, except for the 0.5N/0h group (n = 2) and 0.25N/6h group (n = 4). *p < 0.05; **p < 0.01; ***p < 0.001; NS = not significant. f) Stratification of limbal marginal capillary loop diameters from previous panel e, between male and female mice following alkali injury from 0–6 h at the intermediate dose post-0.250N. Data are presented as mean ± SEM. One-way ANOVA, loop diameter: p ≤ 0.0001, followed by Tukey’s multiple-comparison test; ***p < 0.001, NS = non-significant.

To determine if sex influence these responses, male and female mice were compared. Although both showed significant changes from controls, no differences in loop dilation between sexes was assessed at this stage (**Fig. 3F**). The increase in loop diameters shortly after exposure indicates that vasodilation is an ultra-early and robust event, consistent with acute inflammation hallmarks of vasodilation and increased permeability preceding leukocyte infiltration and tissue repair. Combined, these aligned alterations in vascular caliber with progressive vasodilation can lead to an increase in intravascular hydrostatic pressure and blood flow, promoting transudation of ultrafiltrate of blood plasma and early swelling at the injury site.

### Changes to lymphatic vessels

In the control, fine green LYVE-1 stained and curled lymphatic capillaries are connected by a large vessel encircling the cornea. Immediately after chemical burn, lymphatic vessels extended towards the central cornea (**Fig. 4a**), most evident in the 0.25N treated corneas at 2 and 6 hours. At 0.5N concentration, vessels dilated with more diffuse LYVE-1 staining suggestive of rupture. These findings reveal a rapid, spatially constricted lymphatic response following injury (**Fig. 4a, d**). Extension lengths increased significantly at 0.125N and 0.25N immediately post-injury, suggesting that lymphatic endothelial cells actively project into inflamed corneal area (**Fig. 4a, b**). These narrow, stretched extensions likely represent early blind-ended elongation, giving rise to pre-collectors. In the ultra-early 0.125N group, the extensions lengths significantly peaked and quickly returned to baseline by 2 hours or below by 24 hours (**Fig. 4a**). In contrast, the 0.25N group showed persistent extension which was obvious up to 6 hours (**Fig. 4b**), with no significant change in the 0.5N group (**Fig. 4c**). Interestingly, at 24 hours with 0.25N, thickened collecting arcs appeared compared with earlier times. The absence of extensions in the 0.5N group suggests that excessive injury may hinder or delay lymphatic engagement. These early extensions may facilitate local fluid clearance or immune surveillance, as evidenced especially at 0.125N and 0.25N (**Fig. 4d**). Overall, these “deployments” occurred without area expansion, indicating directional extension rather than early lymphangiogenesis (**Fig. 4e**). Despite variability, limbal lymphatic vessel area coverage remained relatively stable compared to controls, with slight immediate significant increases above 0.25N, and decrease with 0.5N at 24 hours. For a more focused analysis, we compared sexes for these responses. Lymphatic area showed no differences across all groups (**Fig. 4f**), however lymphatic extensions significantly increased in both female and male post-injury versus controls, with males displaying greater extensions compared to females (**Fig. 4g**). Overall, these findings indicate an ultra-early directional extension in both females and males rather than global lymphangiogenesis in the acute window post-injury, with male-biased enhancement in lymphatic engagement.

**Fig. 4.**
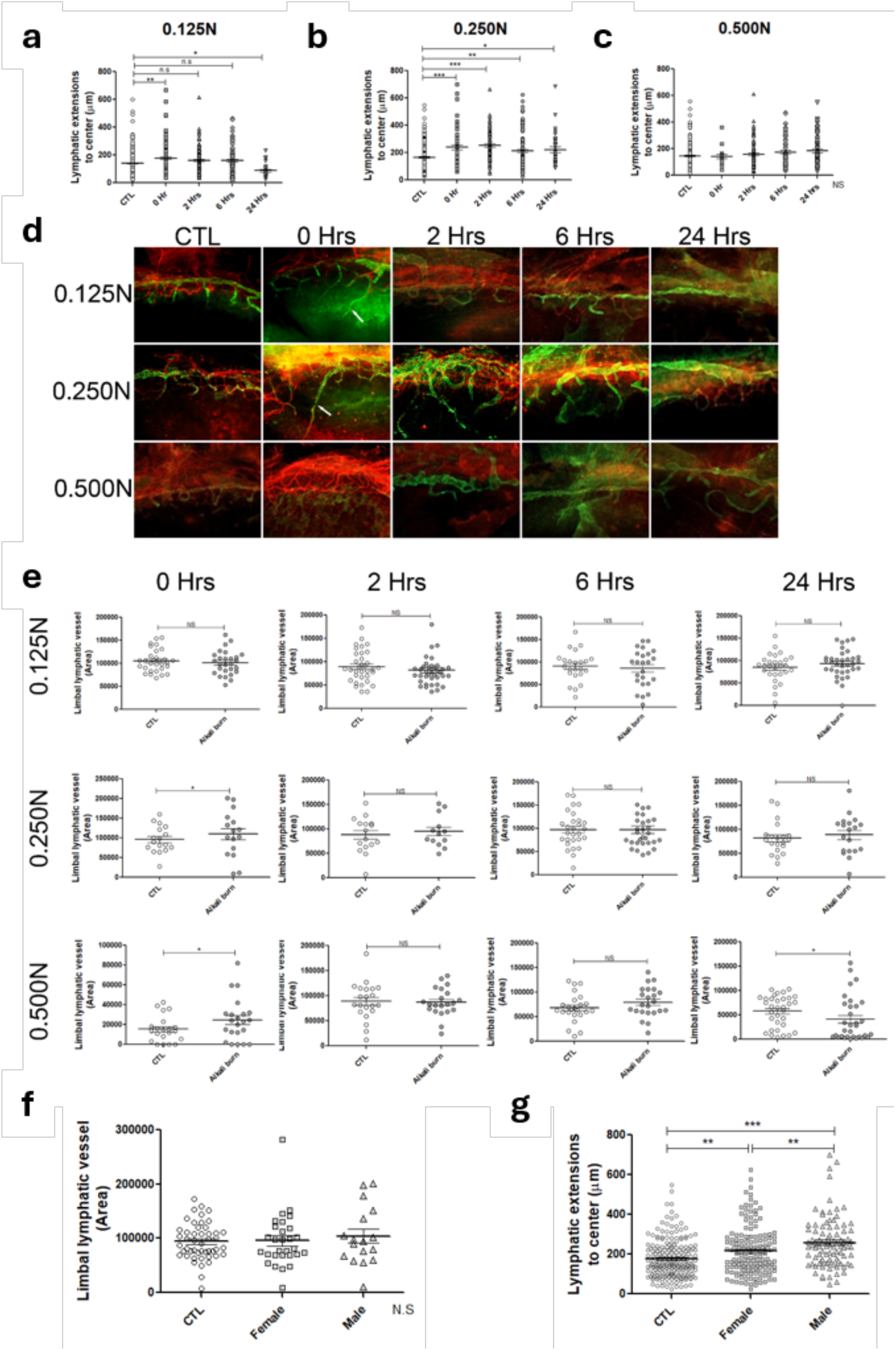
Early lymphatic vascular response in alkali induced acute inflammation. a-c) Quantification of lymphatic vessel extension length (μm) from the corneo-limbal arcade toward the central cornea at 0, 2, 6, and 24 hours post-injury. Multiple extension measurements were obtained across the cornea to characterize spatial variability. One-way ANOVA followed by Tukey’s test was used to compare each time point. ANOVA: a-b) p < 0.0001; c) p = 0.0534. Data represent measurement averaged mean ± SEM. n = 3 mice for most of the analysis, except for the 0.125N/0h group (n = 4), 0.125N/2h group (n = 5), 0.125N/24h group (n = 4), 0.125N/CTL group (n = 16), 0.25N/24h group (n = 4), 0.25N/CTL group (n = 13), and 0.5N/CTL group (n = 12). Asterisks indicate significant post hoc differences: *p < 0.05; **p < 0.01; ***p < 0.001; NS = not significant. d) Representative flatmounts images showing lymphatic (green, LYVE,1) and blood vessels (red, CD31) in control and post-burn corneas. n = 3 mice for most of the analysis, except for the 0.125N/0h group (n = 4), 0.125N/2h group (n = 5), 0.125N/CTL group (n = 15), 0.25N/24h group (n = 4), 0.25N/CTL group (n = 13), and 0.5N/CTL group (n = 12). e) Quantification of total limbal lymphatic vessel area using ImageJ ROI delineation. Student T test was used to compare groups. Data represent mean ± SEM. n = 3-4 per group. *p < 0.05, *p < 0.01; NS = not significant. f,g) Sex-stratified quantification of total lymphatic area and limbal lymphatic extensions shown above for the 0–6 h, 0.25N condition. Data are presented as mean ± SEM. One-way ANOVA, area: N.S; extensions: p ≤ 0.0001, followed by Tukey’s multiple-comparison test was used to assess differences between control and injured corneas *p < 0.05, **p < 0.01, ***p < 0.001; NS = non-significant.

### Microvascular leakage

To determine if the burn induced inflammation was accompanied by vascular permeability, we adapted the Miles assay using intravenous injections of Evans Blue to detect and measure the dye-plasma albumin complex extravasation in the corneal stroma exposed to 0.25N NaOH. There was minimal EBA accumulation in the control corneas. At 2 hours post-exposure, blue dye was observed at the periphery of the cornea (**Fig. 5a**). When viewed under fluorescence, small amounts of EBA leakage (red) were observed immediately after the burn (0 hour). The leakage continued to spread into the periphery of the cornea at 2 and 6 hours post-burn, but by 24 hours, the dye had dissipated.

**Fig. 5.**
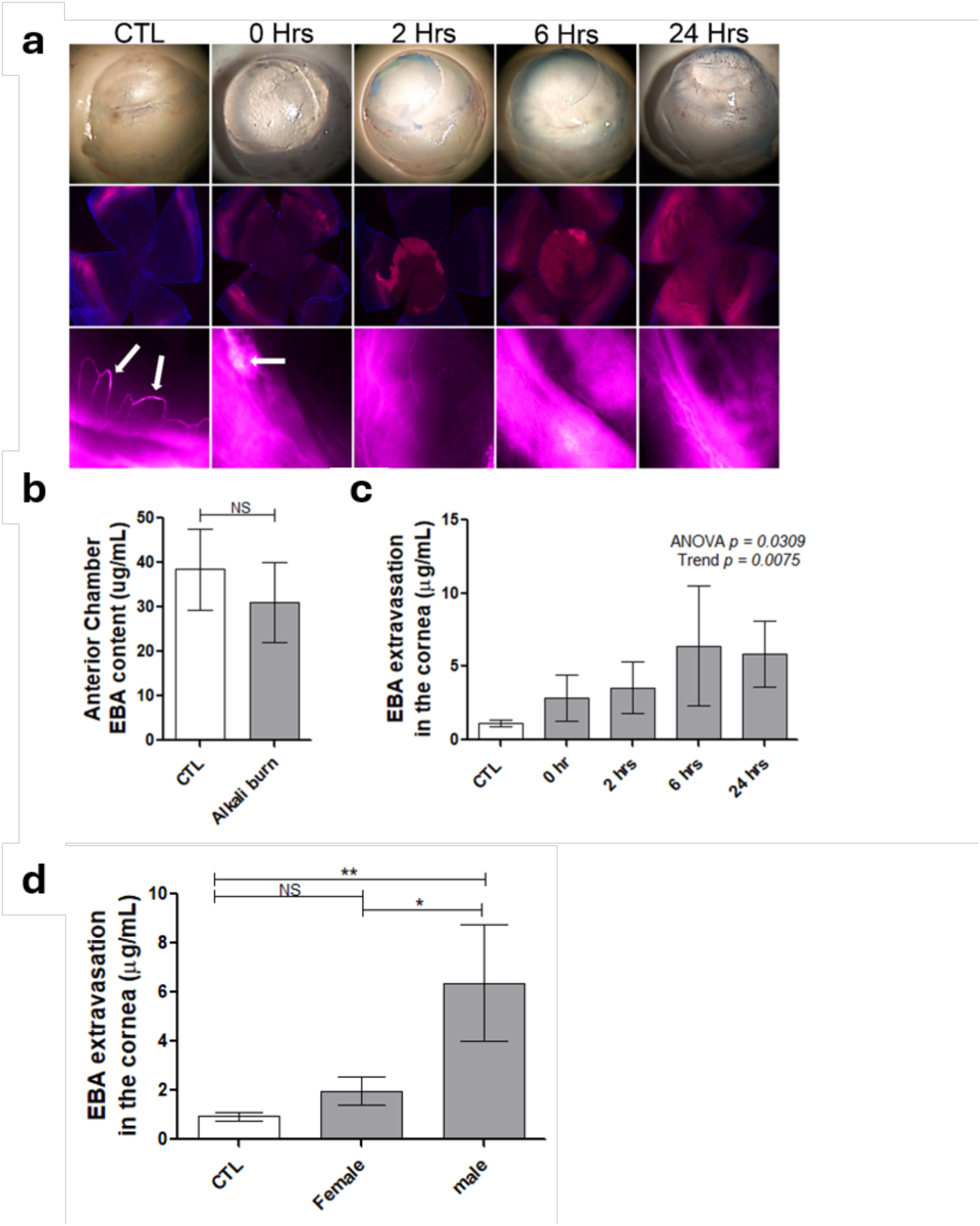
Corneal Vascular permeability following alkali injury. Direct evaluation of corneal vascular permeability following alkali burn. a) Representative macroscopic and fluorescence images of corneas harvested at 0, 2, 6, and 24 hours post-injury following Evans Blue albumin (EBA; seen as a blue dye) injection and 2 hours circulation. White arrows highlight areas of dye extravasation. b) Quantification of EBA content in anterior chamber fluid showing no significant difference between control and injured groups. Student T test was used to compare between groups, presented as mean ± SEM; of n = 8 eyes/group. NS = not significant. c) Fluorometric quantification of EBA in the cornea shows significant dye extravasation immediately and 2 hours post-injury, which plateaus by 6 and 24 hours. One-way ANOVA followed by a post hoc linear trend was used to assess dose–response significance (ANOVA p = 0.0309; trend p = 0.0075). Data are means ± SEM. Replicates for the groups were: 0h group (n = 4), 2h group (n = 6), 6h group (n = 3), 24h group (n = 3), and CTL group (n = 17). d) Sex-stratification of Evans Blue Albumin (EBA) leakage quantification shown in c, from 0–6 h. Measurements were obtained from ipsilateral (OD) injured corneas relative to the contralateral cornea (CTL). Replicates for the groups were: Female (n = 7), male (n = 6), and CTL (n = 13). Data are presented as mean ± SEM. One-way ANOVA, EBA: p = 0.003, followed by Tukey’s multiple-comparison test was used to assess differences between control and injured corneas *p < 0.05, **p < 0.01 NS = non-significant.

In order to confirm that the EBA spread over the cornea was not due to dilution of dye from the aqueous humour, the EBA content of the aqueous chamber of control corneas when compared to that in burnt corneas showed a slight decrease, showing that any EBA spread was not due to dye dilution (**Fig. 5b**). Quantification of EBA extravasation showed a significant rapid increase in dye content right after injury with levels remaining high over 24 hours (**Fig. 5c**). Interestingly, upon gender stratification, vascular permeability significantly increased only in males by approximately six-fold relative controls and notably diverged compared to females (**Fig. 5d**).

### Lymphatic Drainage and Systemic Uptake After Corneal Injury

To investigate the functional engagement of the lymphatic system following central corneal injury, we next assessed limbal lymphatic uptake along structural dynamics within the first half hour post-exposure. These analyses aimed to determine if lymphatic activation coincided with vascular responses. Following topical application of 40 kDa FITC-Dextran to the injured corneal surface, tracer signal was detected at the injury site and tracked over time (**Fig. 6a**). Fluorescent signal progressively spread from the corneal surface over the course of 5 to 20 minutes. At 5 minutes, localized green fluorescence appeared at the limbus in a segmented, slightly radial pattern consistent with initial tracer entry into limbal lymphatic capillaries. The pattern was not diffuse, outlining discrete pathways, suggesting structured vessel uptake rather than non-specific pooling.

**Fig. 6.**
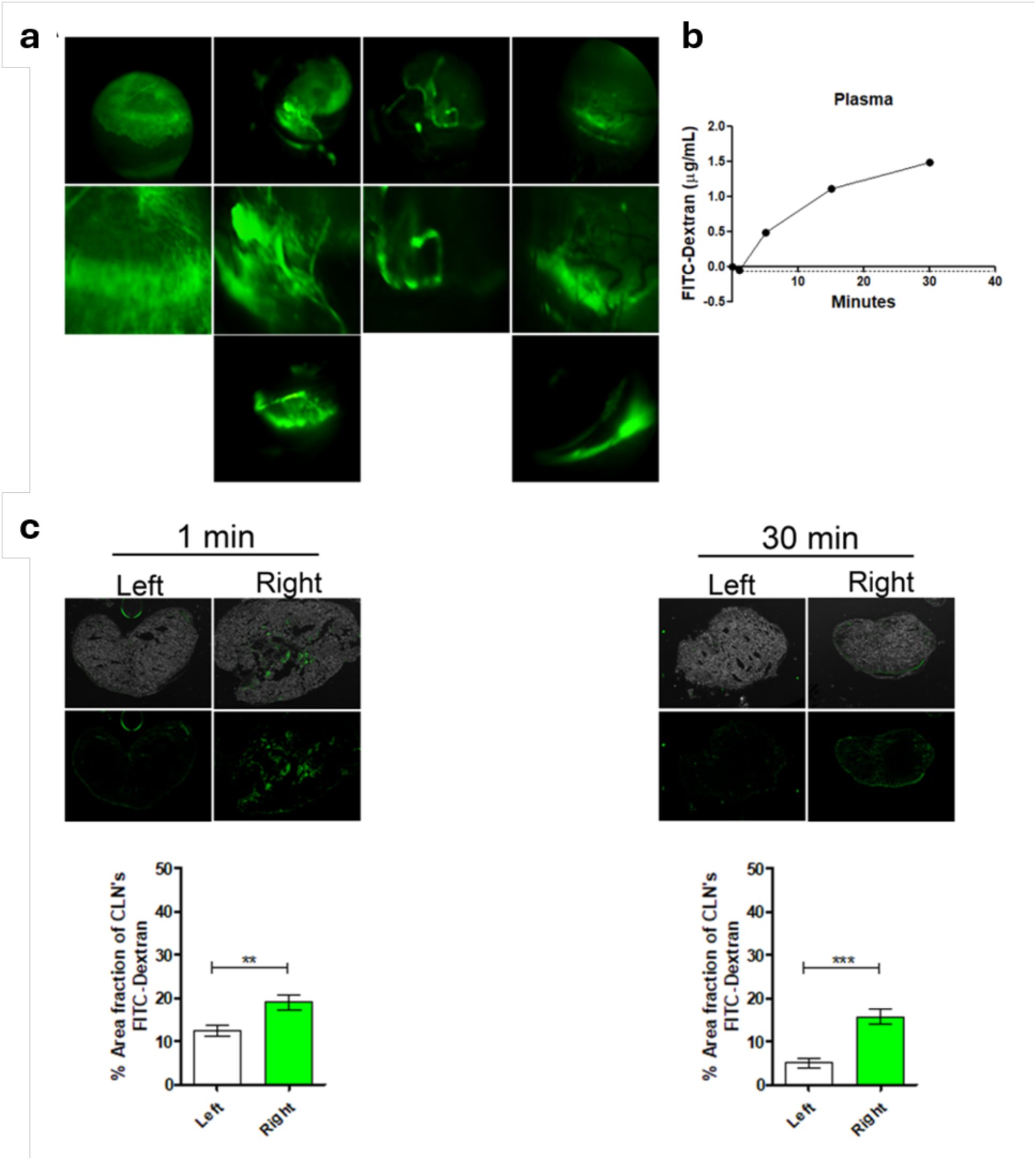
Corneal FITC-Dextran tracer lymphatic uptake following alkali injury. a) Representative live images of the corneal surface acquired using the Micron IV imaging system, showing FITC-Dextran distribution at 5, 10, and 20 minutes post-application in the injured eye compared to the uninjured control (CTL). b) Quantification of FITC-Dextran concentration in plasma over time confirms exclusion of the tracer from systemic circulation. Graph represents average mean of 2 independent experiments, male and female for each time point at 0, 1, 5 and 30 minutes (n = 2), 15 minutes (n = 1; female). c) Representative cryosections of CLNs collected from contralateral (Left) and injured ipsilateral (Right) sides at 1 and 30 minutes post-application n = 2 mice/Time. Green signal indicates FITC-Dextran accumulation in CLNs. Quantification of the percentage area from 5 sectioned images/node covered by fluorescence reveals increased tracer uptake in ipsilateral CLNs over time. Student T test was used to compare groups. Data are presented as mean ± SEM. **, p < 0.01; ***, p < 0.001.

By 10 minutes, the signal intensified following the curvilinear limbal arc, indicating FITC-Dextran was moving through limbal lymphatics, possibly into pre-collector or collector vessels encircling the cornea. At 20 minutes, the signal extended along what appeared to be lymphatic channels, confirming sustained drainage, suggesting that the extended vessels were functionally patent and maintained uptake over time. In contrast, controls showed diffuse signal which indicates injury-induced specific tracer uptakes. FITC-Dextran was also detectable in plasma shortly after topical application, with a marked increase between 5 and 20 minutes post-injury (**Fig. 6b**). This suggests that a portion of the tracer rapidly gained access to systemic circulation, potentially through nasolacrimal drainage (via the conjunctival membrane) or blood and lymphatic pathways activated in response to injury (42, 43).

To assess whether the tracer drained via functional lymphatic pathways, CLNs were analyzed following topical administration (**Fig. 6c**). Consistent with active and time-dependent lymphatic transport from the cornea to regional lymph nodes, quantification of the percentage area covered by FITC-Dextran fluorescence revealed enrichment as early as 1 minute post-application in the injured ipsilateral CLNs (right side) compared to the uninjured contralateral CLNs (left side), which remained relatively stable by 30 minutes, indicating active lymphatic drainage. Nevertheless, a notable signal was detected in contralateral CLNs, suggesting additional physiological clearance route. These findings confirm that newly engaged lymphatic pathways are functionally connected to systemic immune drainage routes within minutes of injury.

### Limited early immune-cell recruitment despite rapid vascular activation

To determine whether the rapid vascular and lymphatic responses observed immediately following alkali injury were accompanied by early inflammatory-cell recruitment, we performed flow cytometric profiling of corneal immune populations immediately (0 h), and at 2, 6 and 24 hours after alkali burn using increasing concentrations of NaOH (0.125N, 0.25N and 0.5N) (**Fig. 7**). Overall, leukocyte recruitment during the first 24 hours was modest compared with the rapid vascular and lymphatic responses described above. Although the total number of CD45⁺ leukocytes increased progressively with injury severity and time, particularly following 0.25N and 0.5N burns, there was little evidence of a rapid, coordinated influx of inflammatory cells during the immediate post-injury period (**Fig. 7a**). Instead, immune-cell accumulation occurred gradually over the subsequent hours, in marked contrast to the more immediate vascular dilation, increased permeability, and lymphatic remodeling.

**Fig. 7.**
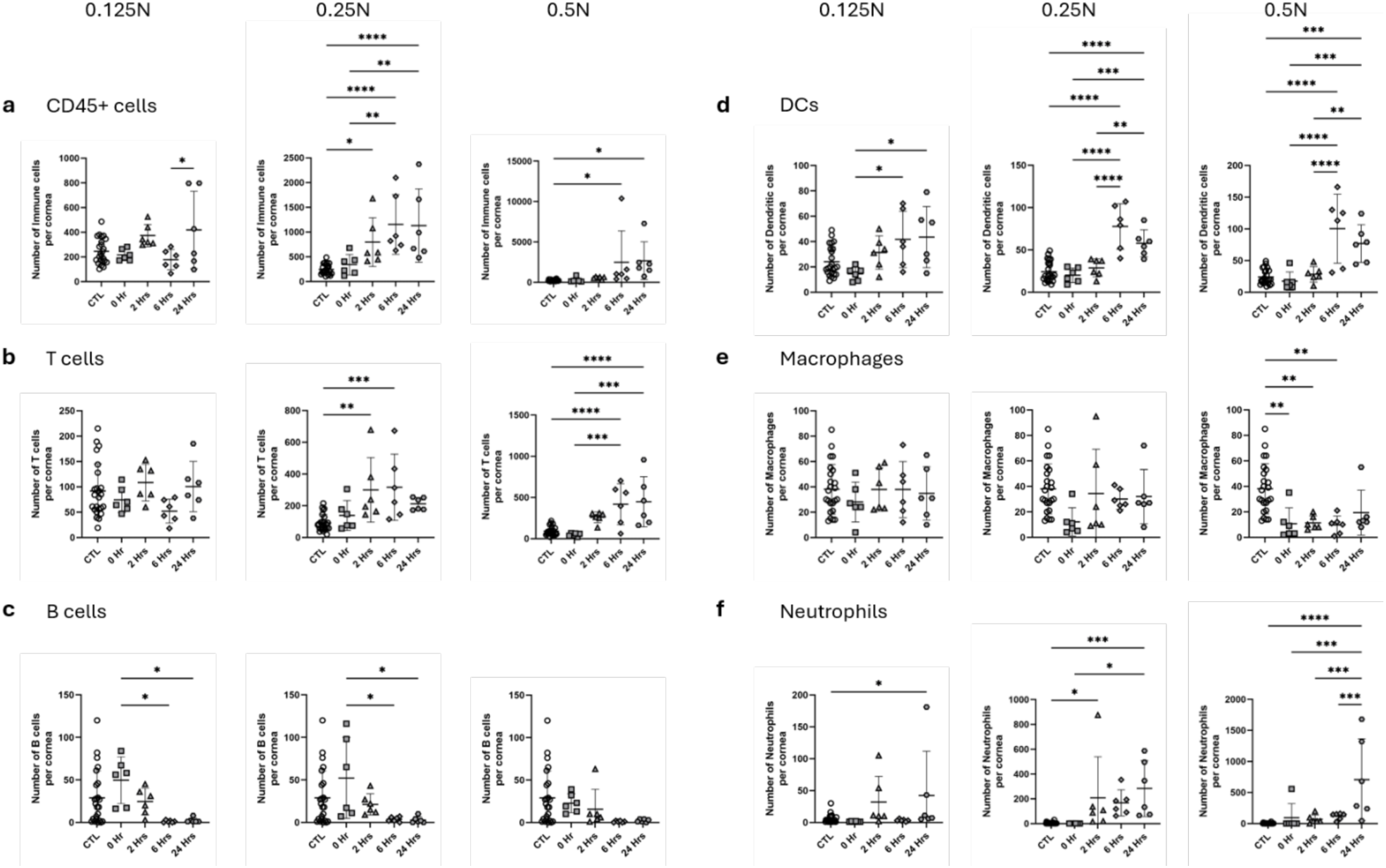
Time course of immune-cell recruitment following corneal alkali injury. Absolute numbers of immune-cell populations per cornea following exposure to increasing concentrations of NaOH (0.125N, 0.25N, and 0.5N) were determined immediately (0 hours) and at 2, 6, and 24 hours after injury and compared with untreated control corneas (CTL). a) CD45+ cells: Total leukocyte infiltration showing force-dependent increases over time, peaking between 6-24 hours. b) T-cells: Infiltration dynamics of total T lymphocytes across force concentrations. c) B cells: Quantitation of B-lymphocyte dynamics per cornea over time. d) Dendritic cells (DCs): Infiltration kinetics of dendritic cell populations. e) Macrophages: Tissue macrophage counts per cornea post-treatment. f) Neutrophils: Acute recruitment kinetics of neutrophils post-injury. Data expression and statistics: Individual data points represent independent biological replicates per experimental group (n = 6), with error bars denoting mean ± SD. Statistical significance was determined using one-way ANOVA with Tukey’s multiple-comparison test (* p<0.05, p<0.01, *** p<0.001, **** p<0.0001).

Analysis of individual immune-cell populations demonstrated a similar pattern. T cells remained relatively stable following mild injury but increased differentially after 0.25N and 0.5N exposure (**Fig. 7b**). No response was seen at 0.125N, while at 0.25N, we saw an increase at 2 and 6 hours; however, the significant relevance was due to a single mouse variant within the pattern. At 0.5N, the response occurred only at 6 and 24 hours. By contrast, B cell numbers generally declined over time, particularly after moderate injury (**Fig. 7c**). Of note, the dendritic cells showed the most consistent increase, rising progressively with all alkali concentrations at 6 and 24 hours after injury (**Fig. 7d**). Macrophage numbers remained relatively low throughout the early response, most evident at later times with 0.5M exposure (**Fig. 7e**), while neutrophils levels, although low showed an increase with 0.25N at 2-24 hours (**Fig. 7f**). Monocytes showed some later recruitment at low and moderate exposure, while few NK cells were recruited (**Supplemental Fig. S3**). The remaining CD45+ immune cells also showed later recruitment at moderate and high doses (**Supplemental Fig. S3**).

Collectively, these findings indicate that leukocyte recruitment during the first 24 hours after alkali injury is modest and delayed relative to the immediate vascular response. Thus, the earliest phase of alkali-induced corneal inflammation is dominated by rapid alterations in limbal blood and lymphatic vessels, with inflammatory-cell recruitment representing a secondary later event.

Given the pronounced sex differences in vascular permeability and lymphatic responses, we next examined whether early immune-cell recruitment also differed between females and males (**Fig. 8**). This included the total number of immune cells defined by CD45 expression (**Fig. 8a**), as well as neutrophils (**Fig. 8b**), monocytes (**Fig. 8c**), macrophages (**Fig. 8d**), dendritic cells (**Fig. 8e**), T cells (**Fig. 8f**), B cells (**Fig. 8g**) and NK cells (**Fig. 8h**). Only modest differences were observed among individual immune-cell populations, such as a decrease in macrophages. However, there was no evidence of a broad leukocyte accumulation. Sex-stratified analysis revealed only limited differences in early immune-cell populations. Although selected differences were observed in monocyte and macrophage numbers, there was no broad sex-dependent increase in leukocyte recruitment during the first 2 hours after injury. The principal exception was a transient increase in monocytes and macrophages in males at 2 hours after injury that was not observed in females. Thus, the more pronounced early vascular responses observed were not accompanied by a correspondingly greater inflammatory-cell influx.

**Fig. 8.**
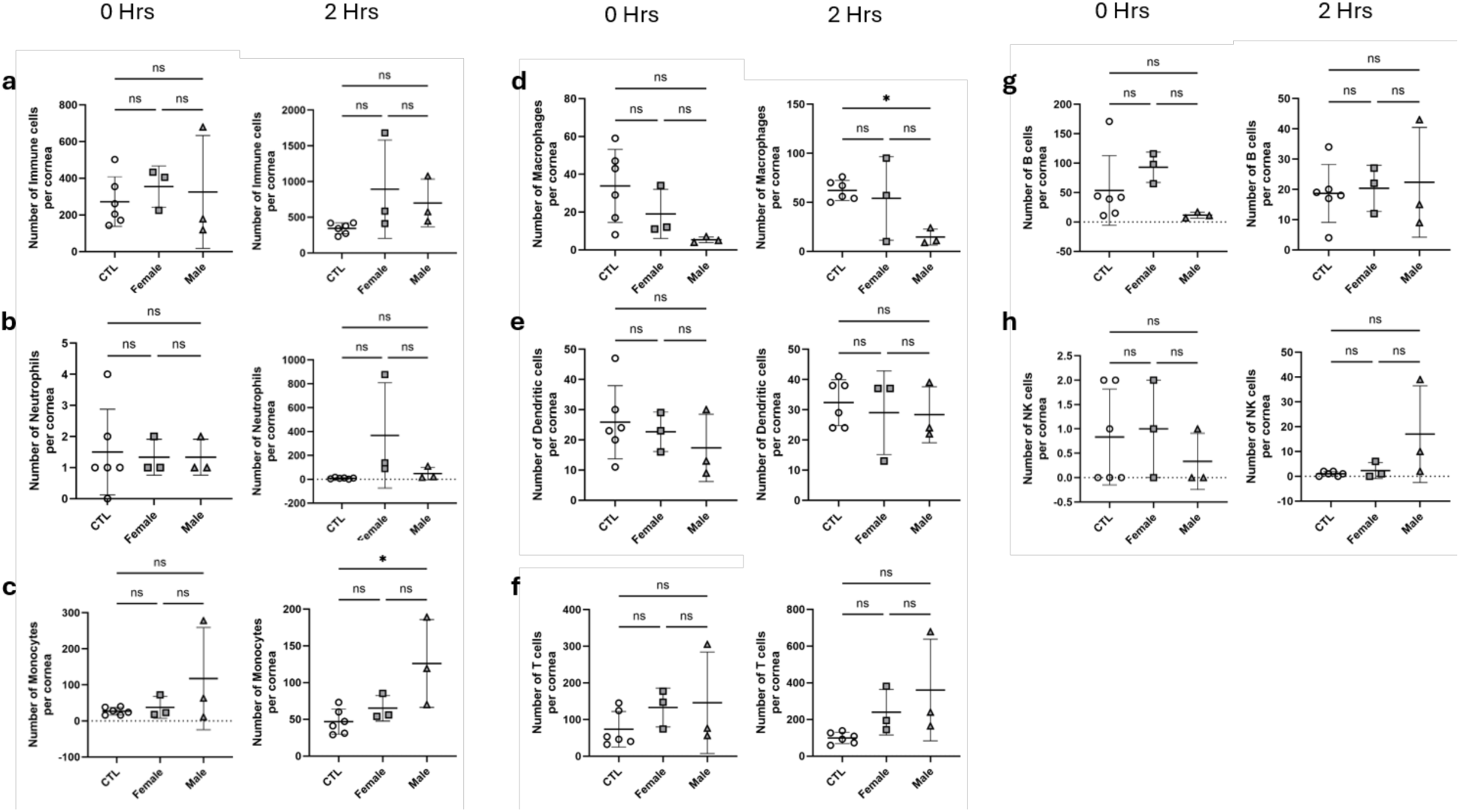
Early corneal immune-cell populations in males versus females following alkali injury. Flow-cytometric quantification of immune-cell populations in the cornea immediately (0 hours) and 2 hours following alkali burn injury. Data are shown for the combined cohort (CTL) and stratified by sex (female and male). a) Total CD45⁺ immune cells, b) neutrophils, c) monocytes, d) macrophages, e) dendritic cells, f) T cells, g) B cells, and h) NK cells per cornea. Each symbol represents an individual cornea (n = 6 for CTL, n = 3 for males and females), while horizontal bars indicate mean ± SD. Statistical comparisons were performed between the indicated groups using one-way ANOVA with Tukey’s multiple-comparison test. *P < 0.05; ns, not significant.

A more detailed longer-term difference between males and females can be found in the supplemental figures (**Supplemental Fig. S4-S12).** A preferential 6 hours recruitment of CD45+ immune cells was observed in males (**Supplemental Fig. S4**). Equal accumulations of T cells at 24 hours post-injury were seen in both sexes (**Supplemental Fig. S5**). A preferential decrease in macrophages at 0.125N at 24 hours was seen in males, while both sexes showed similar decreases at 6 and 24 hours with higher level alkali concentrations (**Supplemental Fig. S8**). Overall, the findings further support the conclusion that the immediate response to corneal alkali injury is dominated by rapid vascular and lymphatic remodeling that precedes inflammatory-cell recruitment.

## IV. Discussion

Acute inflammation typically involves vasodilation, vascular permeability and leukocyte infiltration. However, the immediate engagement of the limbal microvasculature in the normally avascular cornea remains poorly characterized (44). Although inflammatory-cell infiltration has long been considered a hallmark of corneal injury, comparatively little attention has been paid to the vascular events that precede leukocyte recruitment and initiate the inflammatory response. Defining these earliest events is important because they establish the tissue microenvironment that ultimately regulates immune-cell trafficking, tissue repair and pathological remodeling. Our study addresses this gap by characterizing vascular and immune responses within minutes following alkali injury, highlighting the importance of this ultra-early phase in shaping subsequent inflammatory and healing processes.

Using a central alkali burn model in mice, we quantified limbal blood and lymphatic responses during the first 24 hours after injury. Within minutes, limbal arcade capillaries dilated and leaked EBA into the stroma, confirming microvascular permeability. Concurrently, blind-ended lymphatic capillaries extended toward the injury site and functionally took up FITC-labeled dextran. Importantly, these responses occurred without evidence of angiogenesis or increased vascular density within the 24-hour period following central corneal burns with NaOH concentrations up to 0.25N. stromal EBA may reflects active limbal extravasation or passive aqueous humor diffusion. Additionally, EBA in the anterior chamber may reflect secondary leakage from the ciliary body or iris, reaching the limbus via Schlemm’s canal or episcleral pathways. These observations emphasize that rapid limbal vascular changes can occur robustly even in the absence of direct limbal injury.

Our findings align with prior evidence indicating that alkali penetrates the mouse cornea extremely rapidly, likely within seconds, leading to immediate mechanical disruption of the extracellular matrix and initiating complex cascades that mobilize vascular and lymphatic responses (45, 46). While the corneal endothelium remained morphologically intact at concentrations ≤0.25N, early osmotic and inflammatory stress may transiently impair endothelial pump function, contributing to stromal edema independent of overt endothelial injury (47). This functional stress, along with limbal vascular permeability, likely contributes to early edema, reinforcing a model in which the limbal plexus acts as a primary route of plasma extravasation during the ultra-early phase. Classical studies support these early events. Early rabbit studies using ≥ 0.2N NaOH showed central corneal edema sparing the limbus, and lacunae formation, consistent with the dose-dependant findings (48). Early corneal opacification and epithelial sloughing within 1–10 minutes were documented, followed by progressive edema from ∼2 to 48 hours, after which re-epithelialization began and angiogenesis was observed around day 5 (49, 50). Although variations exist due to differences in concentration, exposure time, and species-specific responses, these observations support the notion that diffusible factors, rather than direct limbal damage, can rapidly induce limbal vascular responses. While sVERGFR or thrombospondin- 1 may delay angiogenesis, other fast-acting mediators likely drive vasodilation and permeability (51). Evidence from other models further supports this. In rabbits, lipid hydroperoxides caused limbal vasodilation within 3 hours, though without quantitative assessment (52). In rats, trypan blue leakage from limbal vessels distant from the injury site was observed within 1 hour post-cauterization, and neutrophil entry occurred within 36 minutes (53). Rat suture-induced injury models confirmed early limbal loop dilation without neovessels at 48 hours, and dilation plus permeability increases, including fibrin extravasation and distended lymphatics, were reported within 6 hours after silver nitrate cautery (54, 55). Human AS-OCTA imaging even detected limbal vascular density increases as early as 1 day post-cataract surgery (56).

Interestingly, EBA extravasation in our model occurred within minutes, contrasting with previous reports showing leakage at much later time points, such as 6–48 hours after environmental ocular stress or higher-concentration alkali burns (57, 58). These findings collectively suggest that the limbal vasculature is primed for rapid responses across species and injury types.

We also observed higher vascular permeability in male mice. Although research on sex differences in acute corneal burn repair remains limited, this is consistent with a study by Irani et al. (2019), who reported higher angiogenic signaling (COX-2, VEGF-A, VEGFR-2) in male rats (59). VEGF promotes vascular permeability through junctional protein disruption, though this was not described in the study (60). Results vary across species: men have a higher risk of glaucoma following ocular chemical burn, whereas no sex-related differences were noted in rabbits following alkali burns (61, 62).

Neurovascular interactions likely contribute significantly to this rapid plasma extravasation. Corneal sensory nerves, with their exceptional density and rapid signal propagation, are well-positioned to detect and respond to tissue injury (25). Injury-induced nociceptive signaling can trigger neurogenic vasodilatory reflexes and rapid release of neuropeptides such as substance P and CGRP, increasing vascular permeability within seconds (63–65). Indeed, substance P has been shown to induce EBA extravasation in peripheral microvascular beds within 20 minutes of injection, supporting the role of rapid neuropeptide-driven permeability (66). Mast cells, strategically located around limbal vasculature, may further amplify these responses through release of histamine, prostaglandins, leukotrienes, and VEGFs (67, 68). Mast cell activation has been shown to drive early limbal vasodilation and EBA leakage within 6 hours in fungal keratitis models, suggesting that immune-mediated mechanisms complement neurogenic signaling (24). VEGF expression rises within hours post-injury, supporting early vascular permeability and extracellular matrix remodeling via MMP activity (69). Additionally, activation of lymphatic vessels by VEGF-C or VEGF-D enhances fluid clearance and reduces acute edema (70), highlighting the integrated roles of these mediators in shaping the immediate inflammatory environment.

Lymphatic responses also occurred rapidly in our study. While lymphatic sprouting usually starts days after injury, we observed functional lymphatic extension and dextran-FITC uptake within minutes (71, 72). Alkali-induced extracellular matrix distortion and collagen redistribution likely lower mechanical resistance, facilitating lymphatic extension and fluid clearance (5, 45, 73, 74). These early lymphatics response may help reduce oedema and initiate immune surveillance. Indeed, intravital two-photon microscopy has shown that LYVE-1+ lymphatic vessels can support immune cell transmigration within minutes, consistent with our findings (75). The more pronounced early vascular responses observed were not accompanied by a correspondingly greater inflammatory-cell influx underscoring the unique nature of the alterations in the vasculature.

A notable finding of this study was that the kinetics of immune-cell recruitment differed substantially from those of the vascular response. Whereas limbal blood vessels underwent rapid vasodilation accompanied by increased vascular permeability and lymphatic vessels were seen to rapidly extend toward the site of injury, infiltration by immune cells was delayed. Further, these events depended on the severity of the alkali burn and the immune-cell subset examined. Total CD45⁺ leukocyte numbers showed only modest increases immediately after injury, with progressive accumulation becoming evident primarily following exposure to 0.25N and 0.5N NaOH. Among individual populations, DCs showed the most consistent increase over time, suggesting that antigen-presenting cells are among the earliest leukocytes to respond to tissue injury. In this context, DCs are involved in debris clearance and antigen presentation (Xie et al., 2018; Akbarzadeh et al., 2026). They can also coordinate innate and adaptive responses by transporting antigens through lymphatic vessels to regional lymph nodes.

In our model, neutrophils, which are usually seen as the first inflammatory cells recruited after tissue damage, increased mostly following moderate and severe burns and only became prominent several hours after injury rather than within the first minutes. Monocyte recruitment was similarly delayed and was most evident following low- and intermediate-dose injury, whereas macrophage numbers remained relatively stable or even declined following severe injury, possibly reflecting tissue damage, altered survival, or migration from the injured cornea.

In contrast, adaptive immune populations exhibited comparatively limited early changes. T cell numbers remained largely unchanged following mild injury and increased at later time points after moderate and severe burns. By contrast, B cell numbers generally declined during the first 24 hours, particularly following intermediate-dose injury. NK cells remained at very low numbers throughout the observation period and showed little evidence of rapid recruitment. These findings demonstrate that the earliest inflammatory phase following alkali injury is not characterized by broad leukocyte infiltration but instead by the selective and progressive recruitment of distinct immune-cell populations whose magnitude and kinetics depend on the severity of tissue injury.

An additional finding that distinguishes this study is that increasing alkali concentrations did not simply produce a greater magnitude of inflammation and instead qualitatively distinct inflammatory programs. Mild injury (0.125N NaOH) elicited a rapid but largely transient vascular response but with limited leukocyte recruitment. This suggests activation of mechanisms primarily involved in tissue homeostasis and repair. In contrast, intermediate injury (0.25N NaOH) produced the most coordinated inflammatory response, characterized by sustained vasodilation, increased vascular permeability, persistent lymphatic extension, and progressive recruitment of dendritic cells, neutrophils and monocytes. The most severe injury (0.5N NaOH) did not simply amplify these responses. Instead, it was associated with impaired lymphatic extension, reduced macrophage representation, delayed accumulation of several leukocyte populations, and evidence of greater tissue disruption, suggesting that excessive injury engages distinct pathological mechanisms rather than representing a linear intensification of the responses observed at lower doses. These findings indicate that the severity of alkali exposure determines the magnitude, and depending on the severity of the injury, composition, timing and organization of the inflammatory response. Rather than inducing a continuum of increasing inflammation, mild, moderate and severe alkali burns appeared to activate a distinct inflammatory program will likely impact subsequent tissue repair and pathological remodeling.

Collectively, our findings support a two-stage model of the acute response to corneal alkali injury. The first stage is an ultra-early vascular phase, characterized by rapid vasodilation, vascular permeability and functional lymphatic activation that occurs within minutes of injury and is largely independent of inflammatory-cell infiltration. The second stage is a progressive cellular phase in which the magnitude and composition of leukocyte recruitment are determined by injury severity and ultimately influence tissue repair and pathological remodeling. Future studies should include endothelial specular microscopy, endothelial structural/functional assays, cytokine profiling, and live in vivo immune cell imaging to comprehensively map these early dynamics.

## V. Conclusion

Overall, our findings reveal a previously underappreciated ultra-early inflammatory phase characterized by rapid limbal vasodilation, plasma extravasation, and functional lymphatic extension within minutes. Sex-specific responses were also evident, with greater vascular permeability in males and early immune-cell representation in females. By delineating this critical window and associated sex differences, we provide mechanistic insight into corneal inflammation and identify early vascular, lymphatic, and immune dynamics as potential modulators of injury progression and resolution following chemical injuries.

## Acknowledgments

We thank Frederic Duval and Anne-Marie Aubin from the Flow Cytometry Core Facility of CR-HMR, and Mikhail Sergeev from the Microscopy Core Facility of CR-HMR for their help. We also thank Dr. Hien Thai Tu, Andres Oroya, and Ammar Sleitin for technical help and discussions. M.G was supported by a Collaborative Health Research Project – Canadian Institutes of Health Research #433429, Natural Science and Engineering Research Council (NSERC) # CHRPJ 549666 – 20 (MG, CB, MCR) and NSERC Canada, #RGPIN-2017- 05410 (MG). MG also acknowledges support from her Canada Research Chair Tier 1, #950-232398 and the Caroline Foundation Research Chair for Cellular Therapy in the Eye. C.E.R. was supported by the Canadian Institutes of Health Research Foundation grant (159912) (Pl) and National Institutes of Health grant RO1 AI049466 (co-Pl). CER holds a majority stake in lmmunAb Res Inc (Canada).

## Author contributions

Experiments, acquisition and analysis of data: NA and MG: Writing, NA, MG, MG and CER; Experimental assistance: MJL, GL, and Delali SD: Conceptualization and design of the work, and project funding: MCR, BL, CER, and MG.

## Data Availability

All data presented within this manuscript is original. All data that supports the findings of this study are available from the corresponding author upon request. Other data sharing is not applicable as data for which a community-recognized, structured repository exists (e.g. next-generation sequencing data) were not created or analyzed in this study.

## Additional information

The authors respect ethical guidelines for animal welfare. All mice were housed in the Maisonneuve-Rosemont Hospital Research Centre (CR-HMR) animal facility under a 12-hour light-dark cycle with ad libitum access to food and water. The animal care program is accredited with the Certificate of Good Animal Practice (GAP) issued by the Canadian Council on Animal Care (CCAC). All experiments involved in the use of animals were approved, reviewed, and overseen by the institutional Committee for the Protection of Animals (CPA).

