## Supplemental Figures for "Rapid Vascular Activation Precedes Immune Cell Infiltration Following Corneal Alkali Burn": Akla et al. 2026 Supplemental Material.pdf

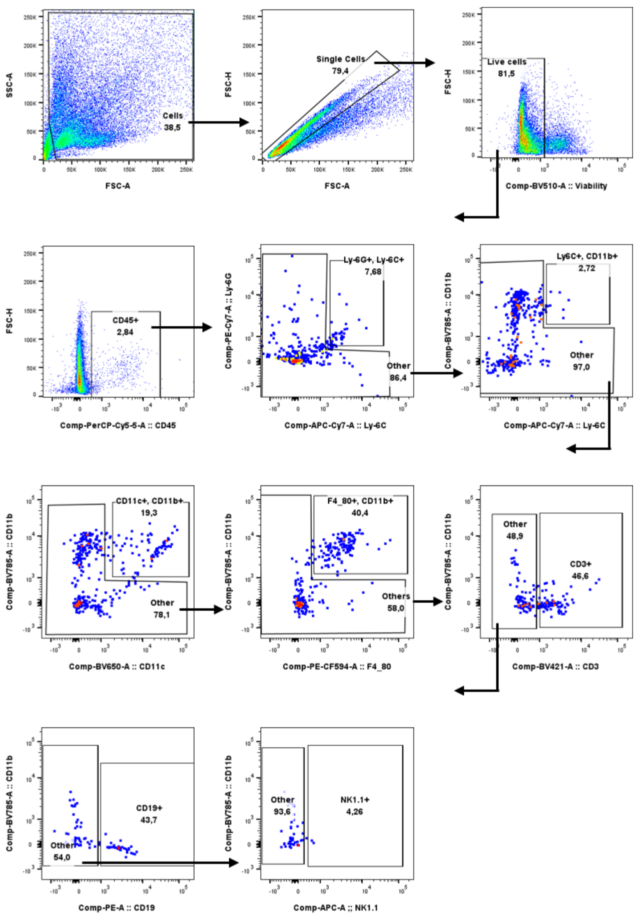


**Supplementary Figure S1**. Flow cytometric gating strategy for identification of immune cell populations in control corneas. Representative flow cytometry plots illustrating the sequential gating strategy used to identify immune cell populations in uninjured control corneas. Following exclusion of debris based on forward- and side-scatter properties, single cells were selected and dead cells excluded using a viability dye. $CD45^+$ leukocytes were then gated and subdivided into $Ly6G^+Ly6C^+$ neutrophils, $Ly6C^+CD11b^+$ monocytes, $CD11c^+CD11b^+$ dendritic cells, $F4/80^+CD11b^+$ macrophages, $CD3^+$ T cells, $CD19^+$ B cells, and $NK1.1^+$ natural killer cells. This gating strategy was applied consistently to all samples analyzed throughout the study.

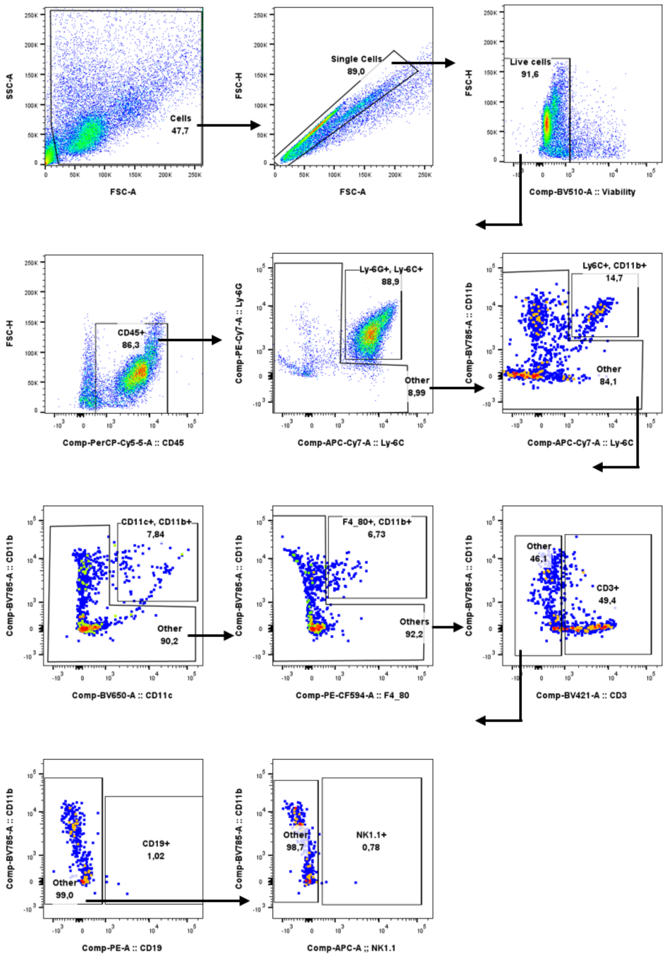


**Supplementary Figure S2.** Flow cytometric gating strategy for identification of immune cell populations in inflamed corneas. Representative flow cytometry plots illustrating the sequential gating strategy used to identify immune cell populations in corneas following alkali injury. The example shown is from a male mouse 24 hours after 0.500 N NaOH injury. Cells were first gated based on forward- and side-scatter properties to exclude debris, followed by selection of single cells and exclusion of dead cells using a viability dye. $CD45^+$ leukocytes were then identified and further subdivided into $Ly6G^+Ly6C^+$ neutrophils, $Ly6C^+CD11b^+$ monocytes, $CD11c^+CD11b^+$ dendritic cells, $F4/80^+CD11b^+$ macrophages, $CD3^+$ T cells, $CD19^+$ B cells, and $NK1.1^+$ natural killer cells. The same gating strategy was applied uniformly to all inflamed corneal samples analyzed in this study.

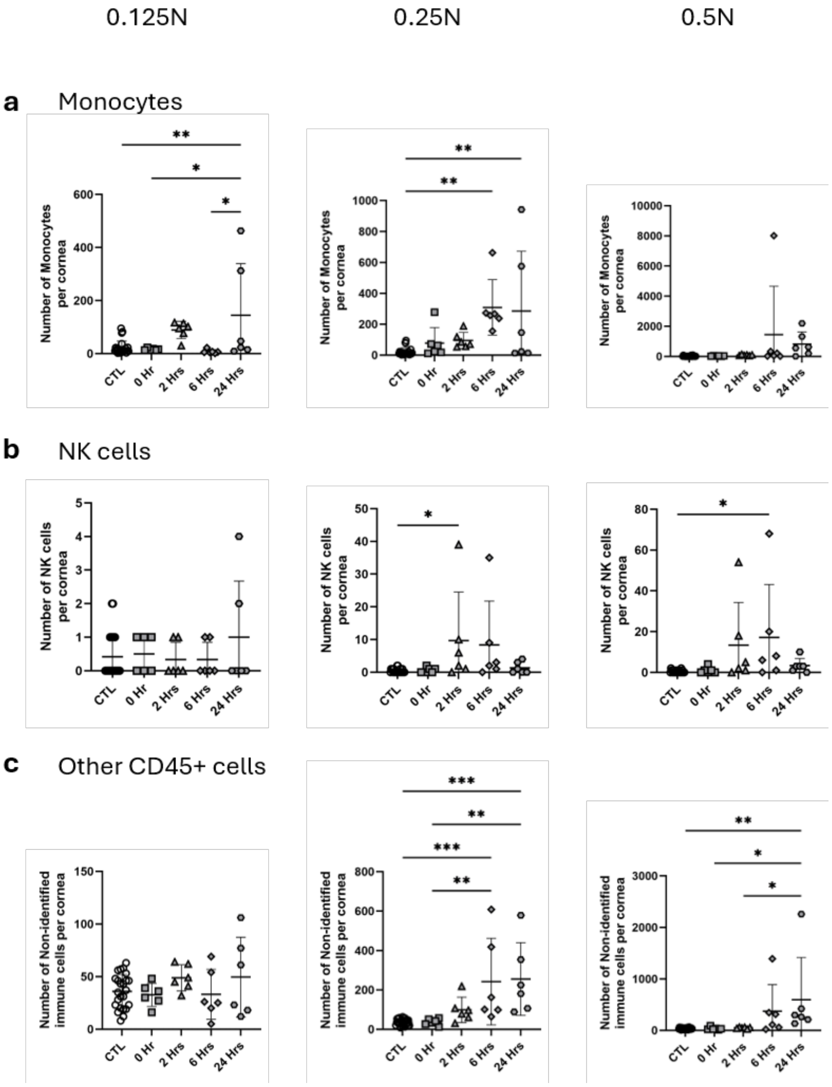


**Supplementary Figure S3**. Monocyte, NK cell, and residual $CD45^+$ leukocyte responses following corneal alkali burn. Flow cytometric quantification of (a) monocytes, (b) NK cells and (c) residual $CD45^+$ leukocytes that were not classified within the major immune-cell populations following corneal alkali injury. Corneas were collected immediately (0 hours), and at 2, 6 and 24 hours after exposure to 0.125N, 0.25N or 0.5N NaOH. The untreated contralateral cornea (CTL) served as the control. Each symbol represents an individual cornea, and horizontal bars indicate the mean ± SEM. Statistical analyses were performed using one-way ANOVA followed by Tukey's multiple-comparisons test. *P < 0.05, **P < 0.01, ***P < 0.001; ns, not significant.

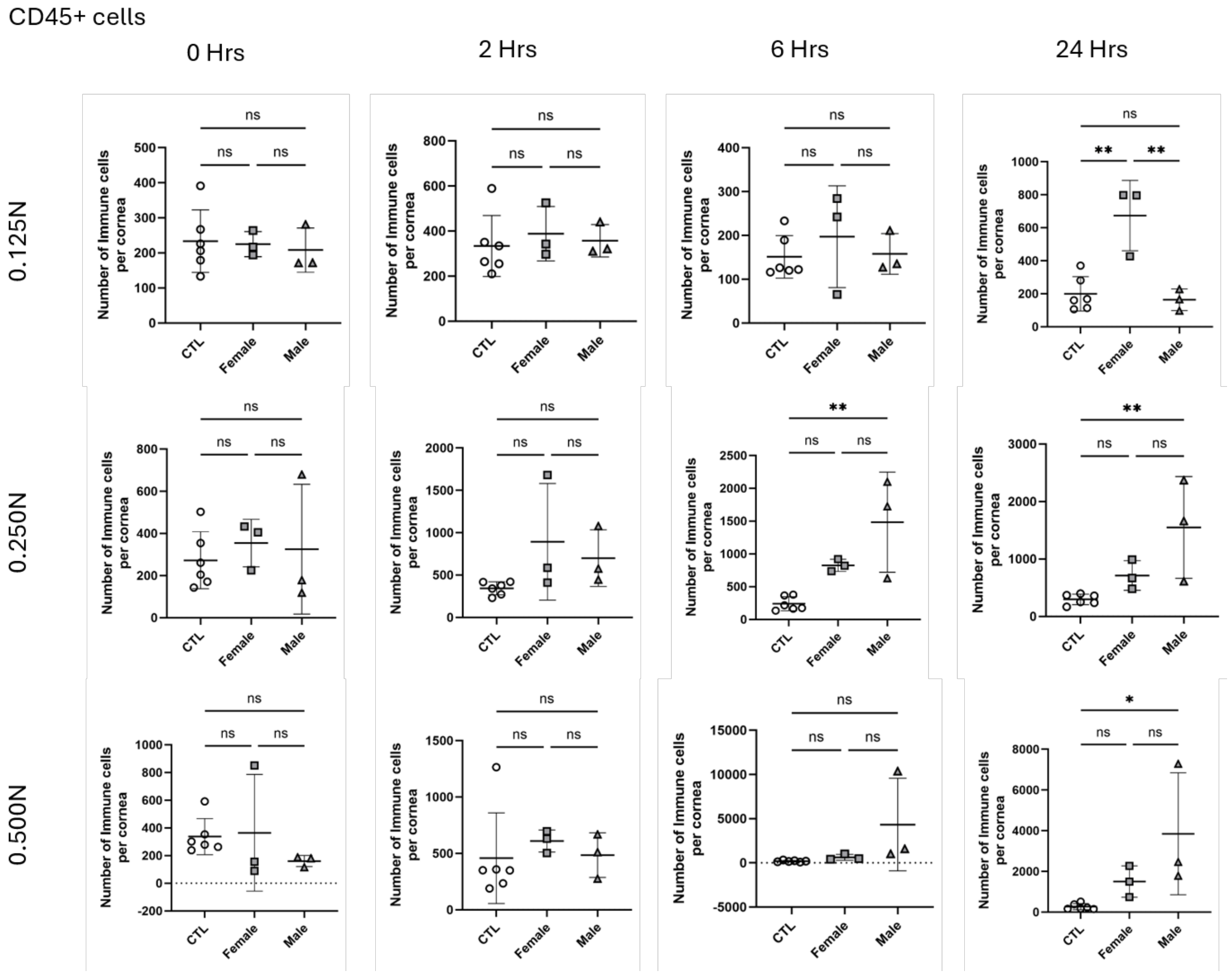


**Supplementary Figure S4**: Quantification of CD45+ cells in male and females following corneal alkali burn. Flow cytometric analysis of CD45+ cells in corneas following alkali injury. Corneas were collected immediately (0 hours), and at 2, 6 and 24 hours after exposure to 0.125N, 0.25N or 0.5N NaOH. Data are shown separately for female and male mice and compared with untreated control (CTL) corneas. Each symbol represents one cornea, and bars indicate mean ± SEM. Statistical analyses were performed using one-way ANOVA followed by Tukey's multiple-comparisons test. *P < 0.05; ns, not significant.

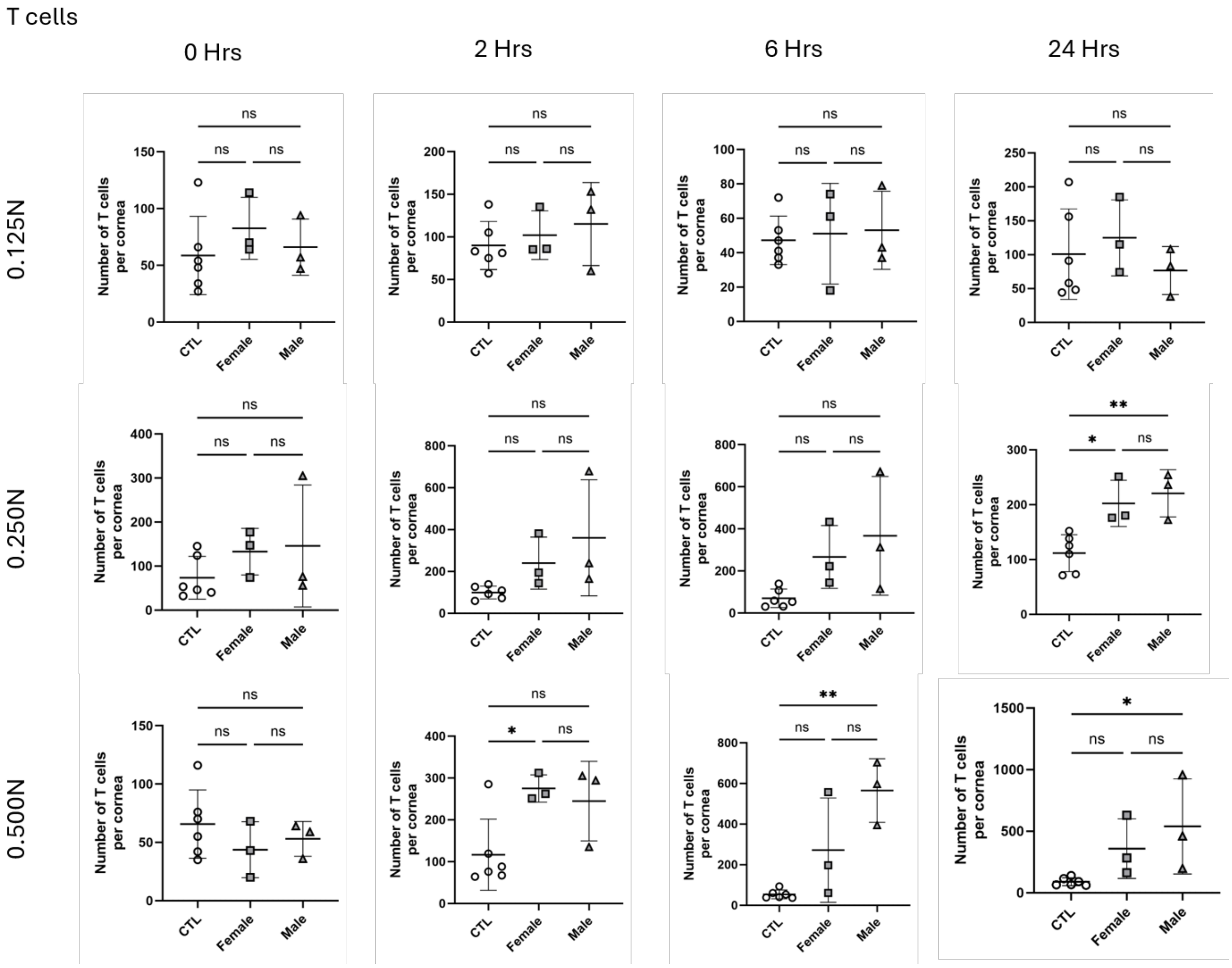


**Supplementary Figure S5:** Quantification of T cells in male and females following corneal alkali burn. Flow cytometric analysis of T cells in corneas following alkali injury. Corneas were collected immediately (0 hours), and at 2, 6 and 24 hours after exposure to 0.125N, 0.25N or 0.5N NaOH. Data are shown separately for female and male mice and compared with untreated control (CTL) corneas. Each symbol represents one cornea, and bars indicate mean ± SEM. Statistical analyses were performed using one-way ANOVA followed by Tukey's multiple-comparisons test. $^{*}P < 0.05$; ns, not significant.

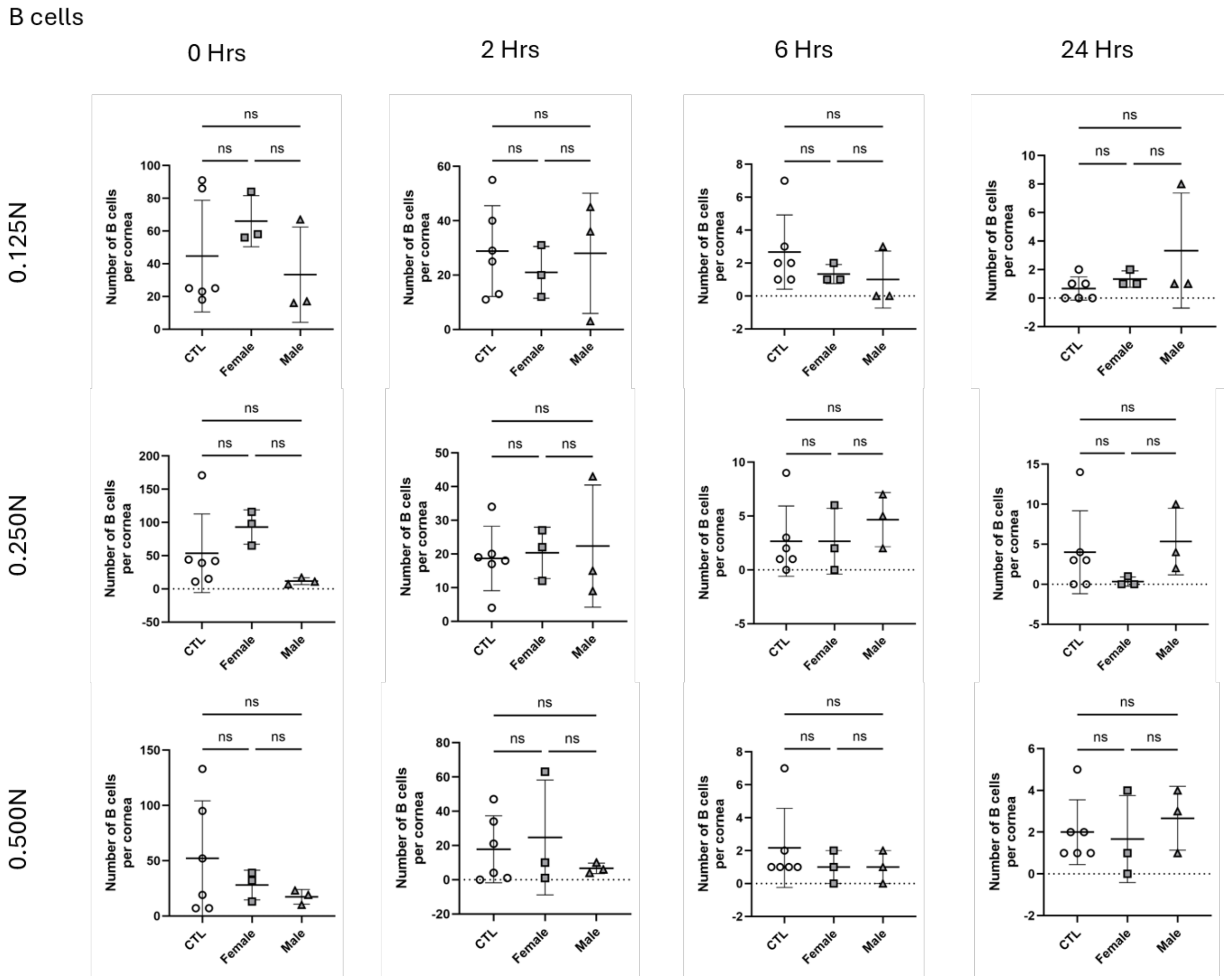


**Supplementary Figure S6:** Quantification of B cells in male and females following corneal alkali burn. Flow cytometric analysis of B cells in corneas following alkali injury. Corneas were collected immediately (0 hours), and at 2, 6 and 24 hours after exposure to 0.125N, 0.25N or 0.5N NaOH. Data are shown separately for female and male mice and compared with untreated control (CTL) corneas. Each symbol represents one cornea, and bars indicate mean ± SEM. Statistical analyses were performed using one-way ANOVA followed by Tukey's multiple-comparisons test. *P < 0.05; ns, not significant.

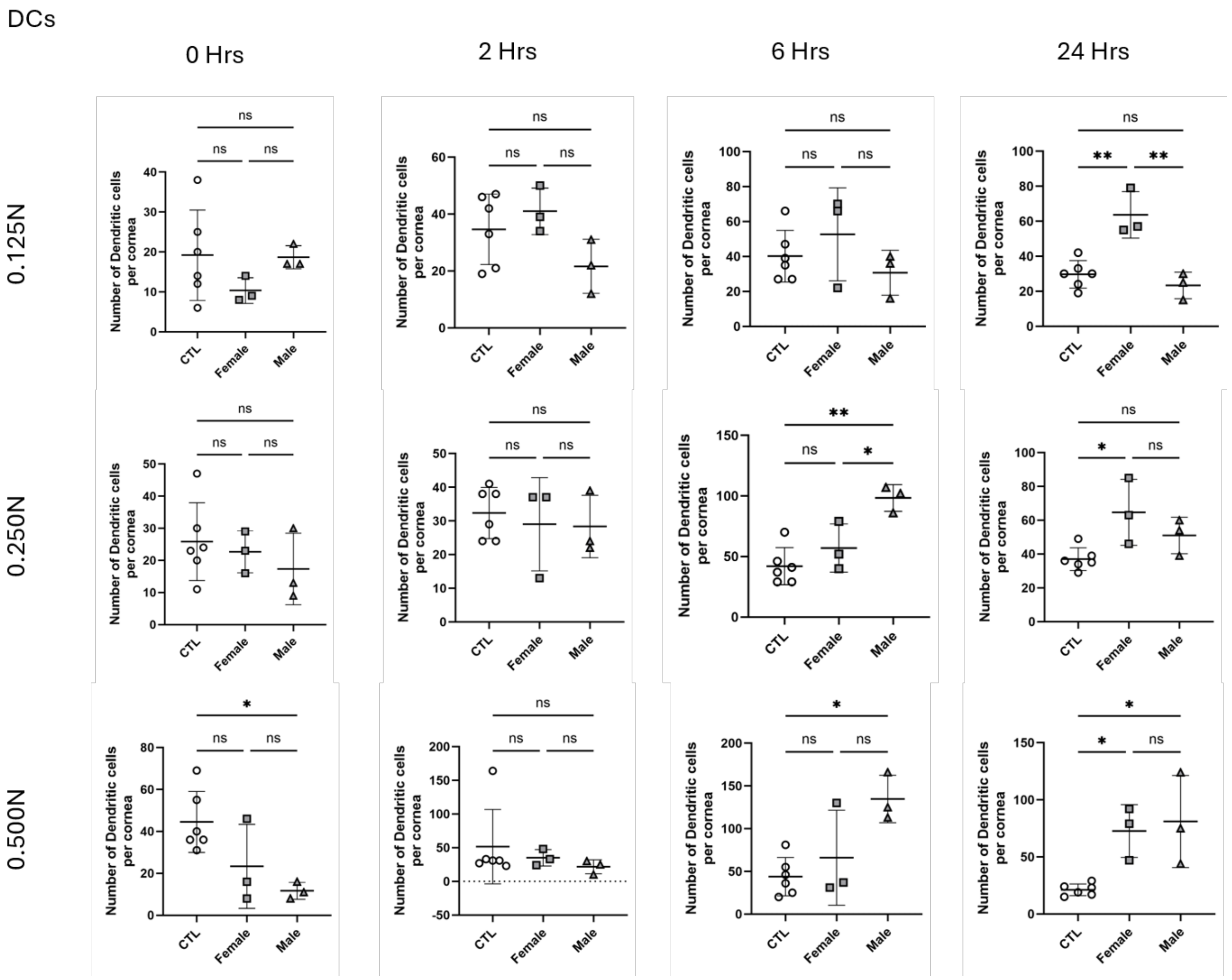


**Supplementary Figure S7:** Quantification of dendritic cells in male and females following corneal alkali burn. Flow cytometric analysis of dendritic cells in corneas following alkali injury. Corneas were collected immediately (0 hours), and at 2, 6 and 24 hours after exposure to 0.125N, 0.25N or 0.5N NaOH. Data are shown separately for female and male mice and compared with untreated control (CTL) corneas. Each symbol represents one cornea, and bars indicate mean ± SEM. Statistical analyses were performed using one-way ANOVA followed by Tukey's multiple-comparisons test. $^*P < 0.05$; ns, not significant.

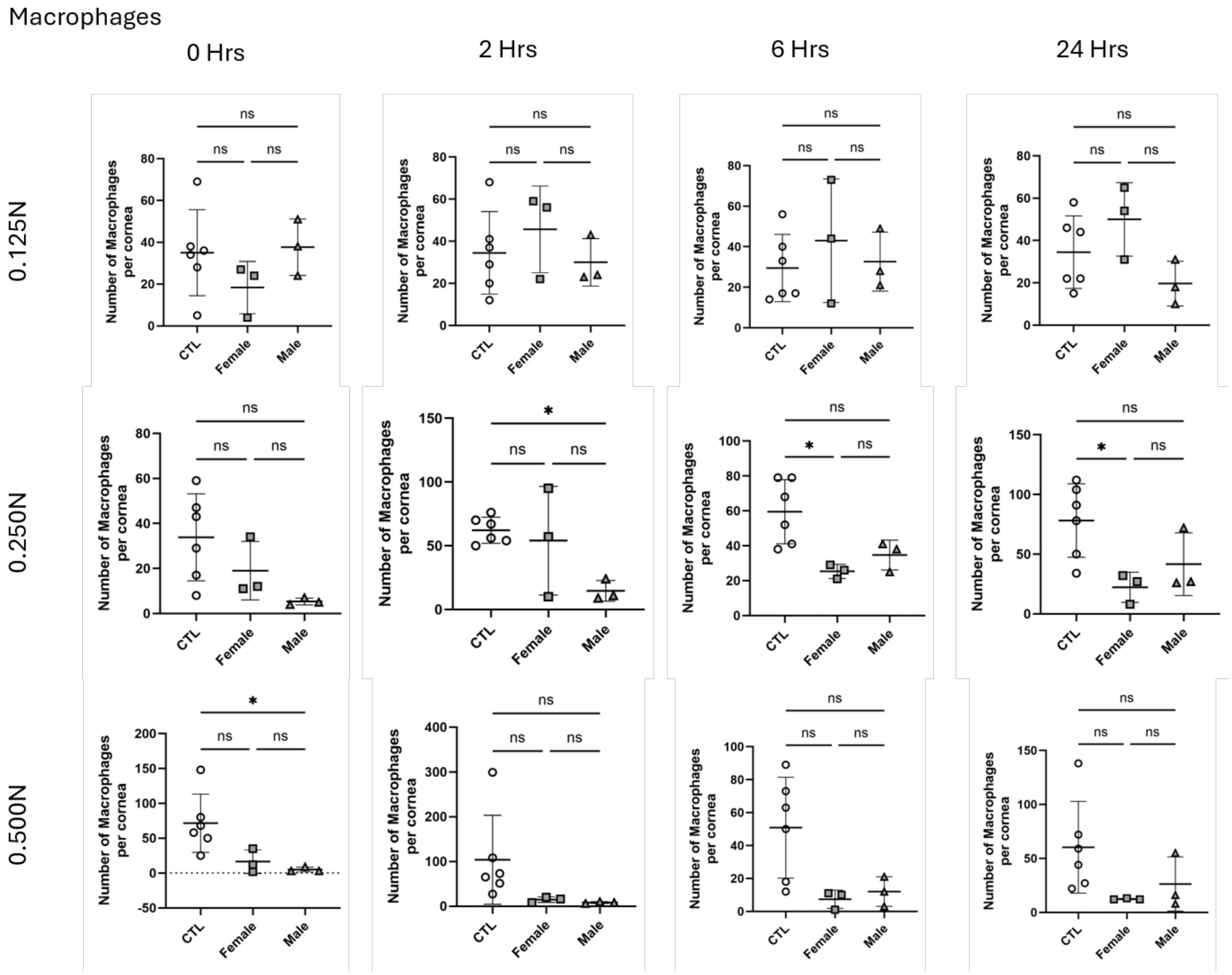


**Supplementary Figure S8:** Quantification of macrophages in male and females following corneal alkali burn. Flow cytometric analysis of macrophages in corneas following alkali injury. Corneas were collected immediately (0 hours), and at 2, 6 and 24 hours after exposure to 0.125N, 0.25N or 0.5N NaOH. Data are shown separately for female and male mice and compared with untreated control (CTL) corneas. Each symbol represents one cornea, and bars indicate mean ± SEM. Statistical analyses were performed using one-way ANOVA followed by Tukey's multiple-comparisons test. *P < 0.05; ns, not significant.

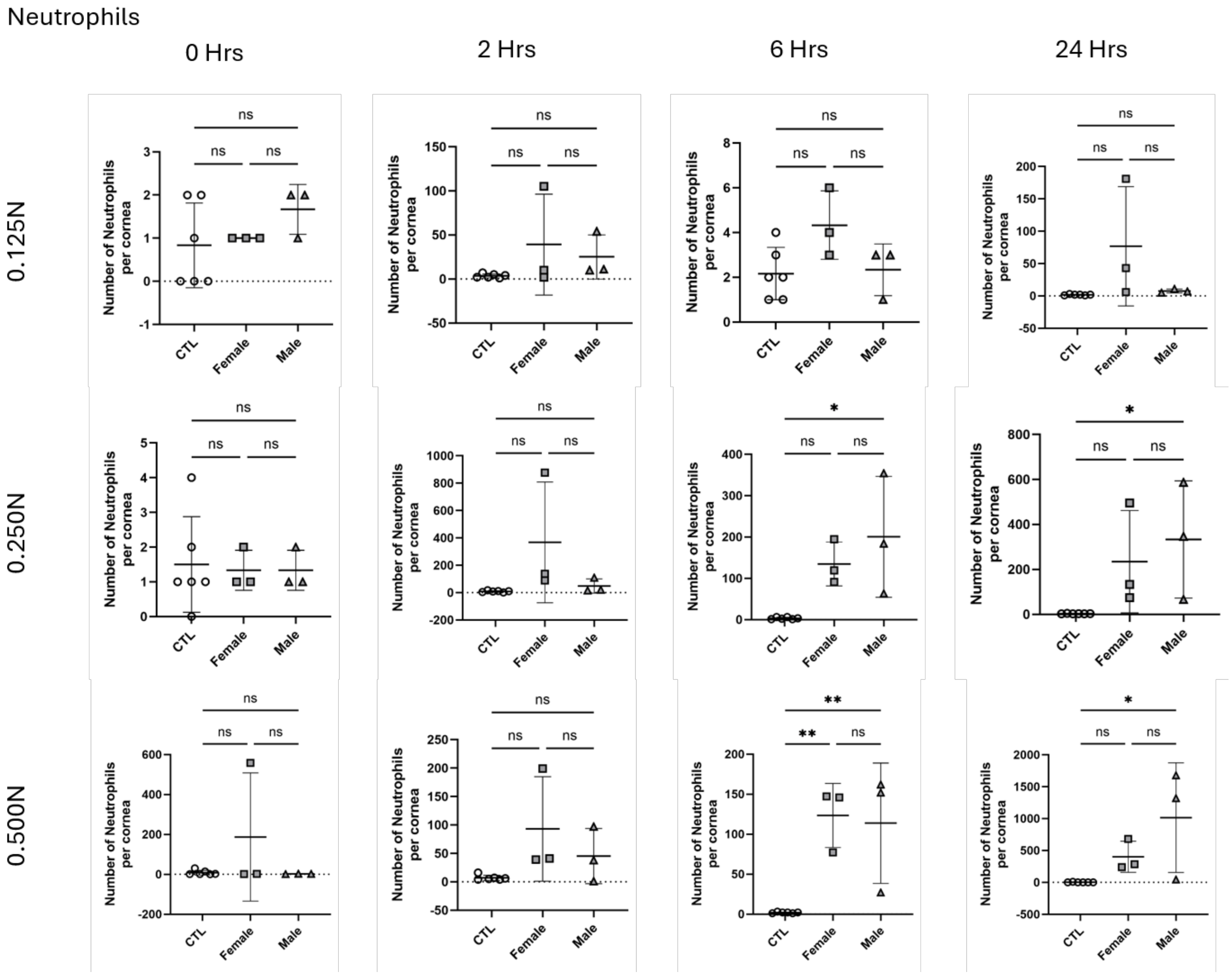


**Supplementary Figure S9.** Quantification of neutrophils in males and females following corneal alkali burn. Flow cytometric analysis of neutrophils in corneas following alkali injury. Corneas were collected immediately (0 hours), and at 2, 6 and 24 hours after exposure to 0.125N, 0.25N or 0.5N NaOH. Data are shown separately for female and male mice and compared with untreated control (CTL) corneas. Each symbol represents one cornea, and bars indicate mean ± SEM. Statistical analyses were performed using one-way ANOVA followed by Tukey's multiple-comparisons test. *P < 0.05; ns, not significant

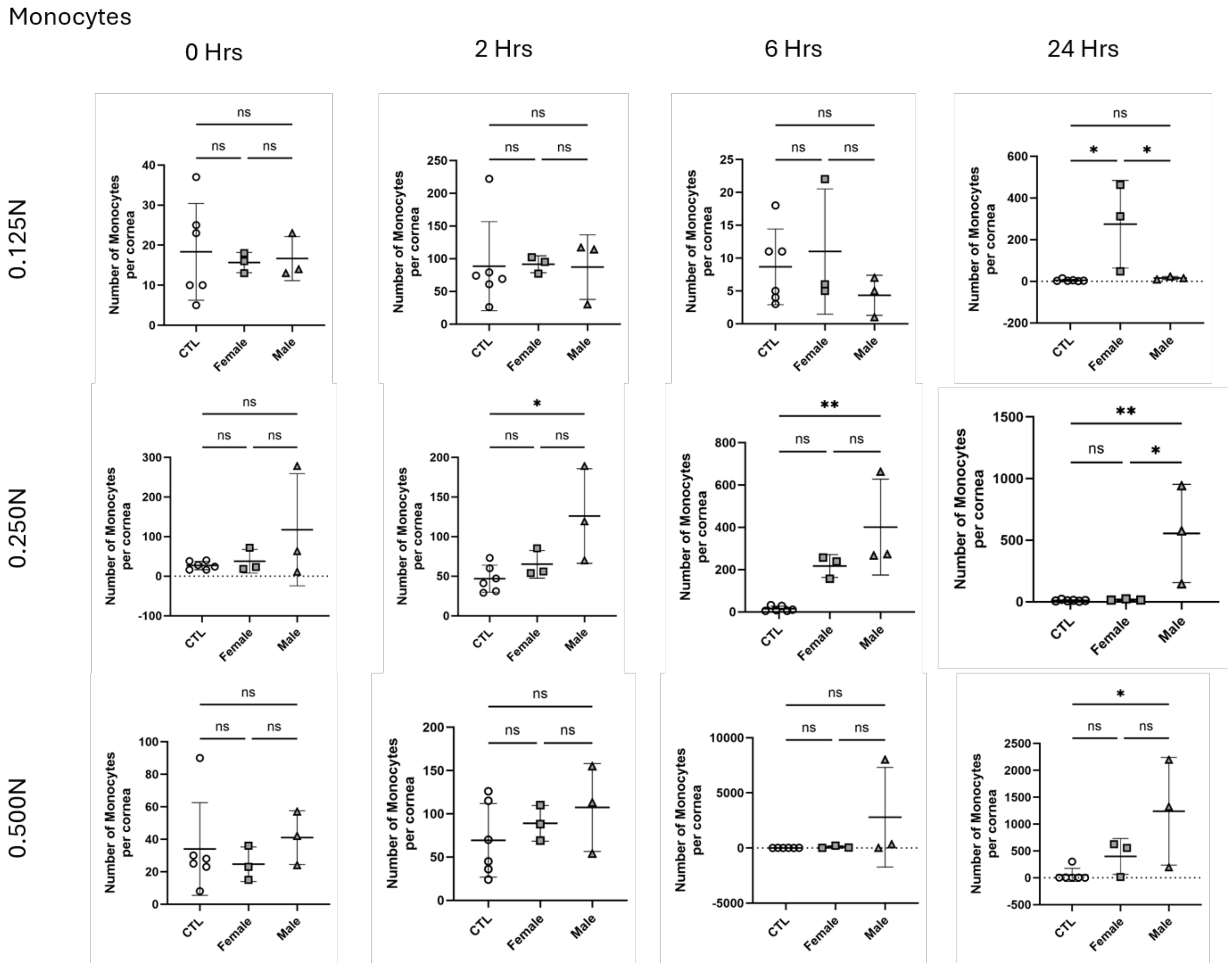


**Supplementary Figure S10.** Quantification of monocytes in males and females following corneal alkali burn. Flow cytometric analysis of monocytes in corneas following alkali injury. Corneas were collected immediately (0 hours), and at 2, 6 and 24 hours after exposure to 0.125N, 0.25N or 0.5N NaOH. Data are shown separately for female and male mice and compared with untreated control (CTL) corneas. Each symbol represents one cornea, and bars indicate mean ± SEM. Statistical analyses were performed using one-way ANOVA followed by Tukey's multiple-comparisons test. *P < 0.05; ns, not significant.

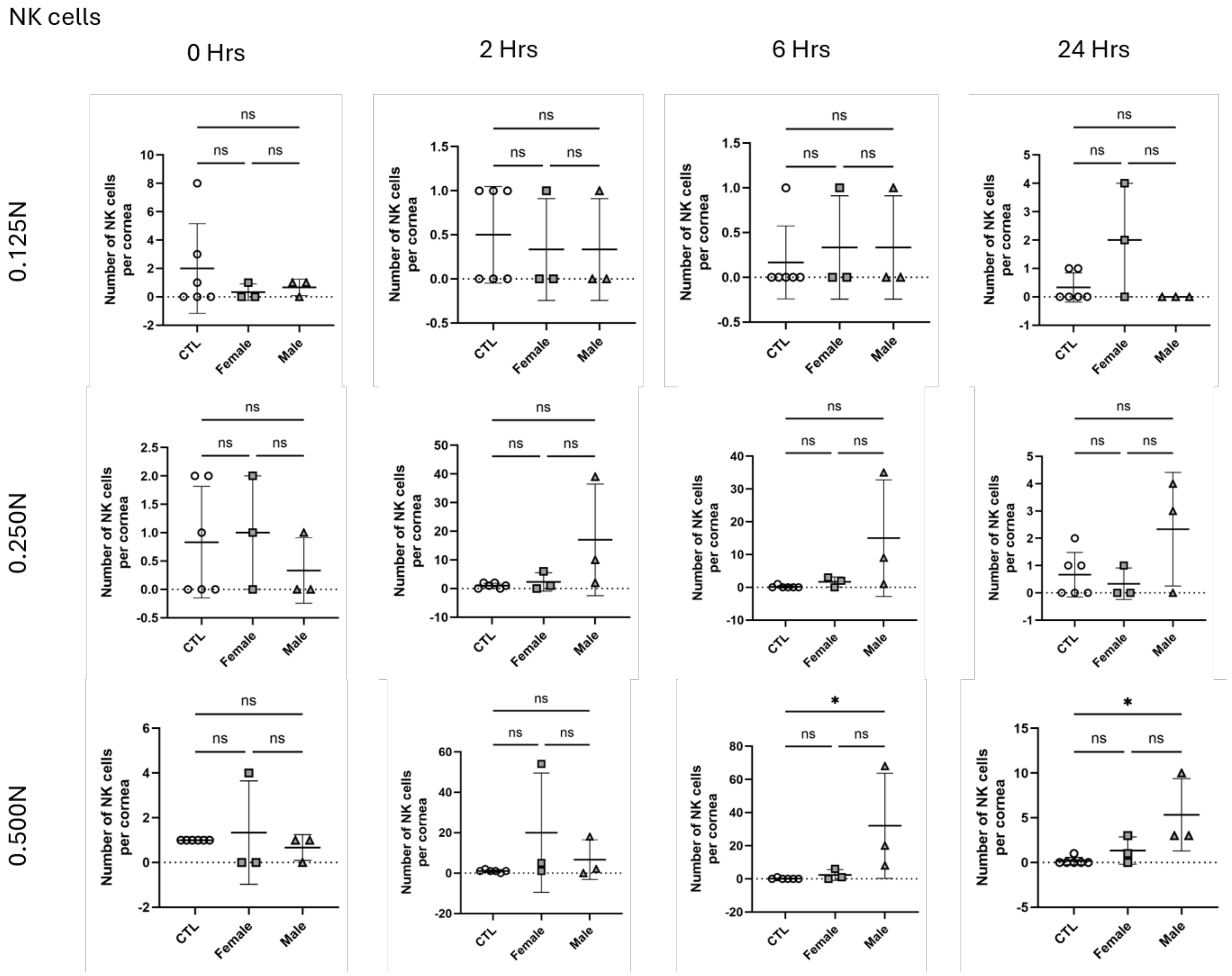


**Supplementary Figure S11:** Quantification of NK cells in males and females following corneal alkali burn. Flow cytometric analysis of NK cells in corneas following alkali injury. Corneas were collected immediately (0 hours), and at 2, 6 and 24 hours after exposure to 0.125N, 0.25N or 0.5N NaOH. Data are shown separately for female and male mice and compared with untreated control (CTL) corneas. Each symbol represents one cornea, and bars indicate mean ± SEM. Statistical analyses were performed using one-way ANOVA followed by Tukey's multiple-comparisons test. *P < 0.05; ns, not significant

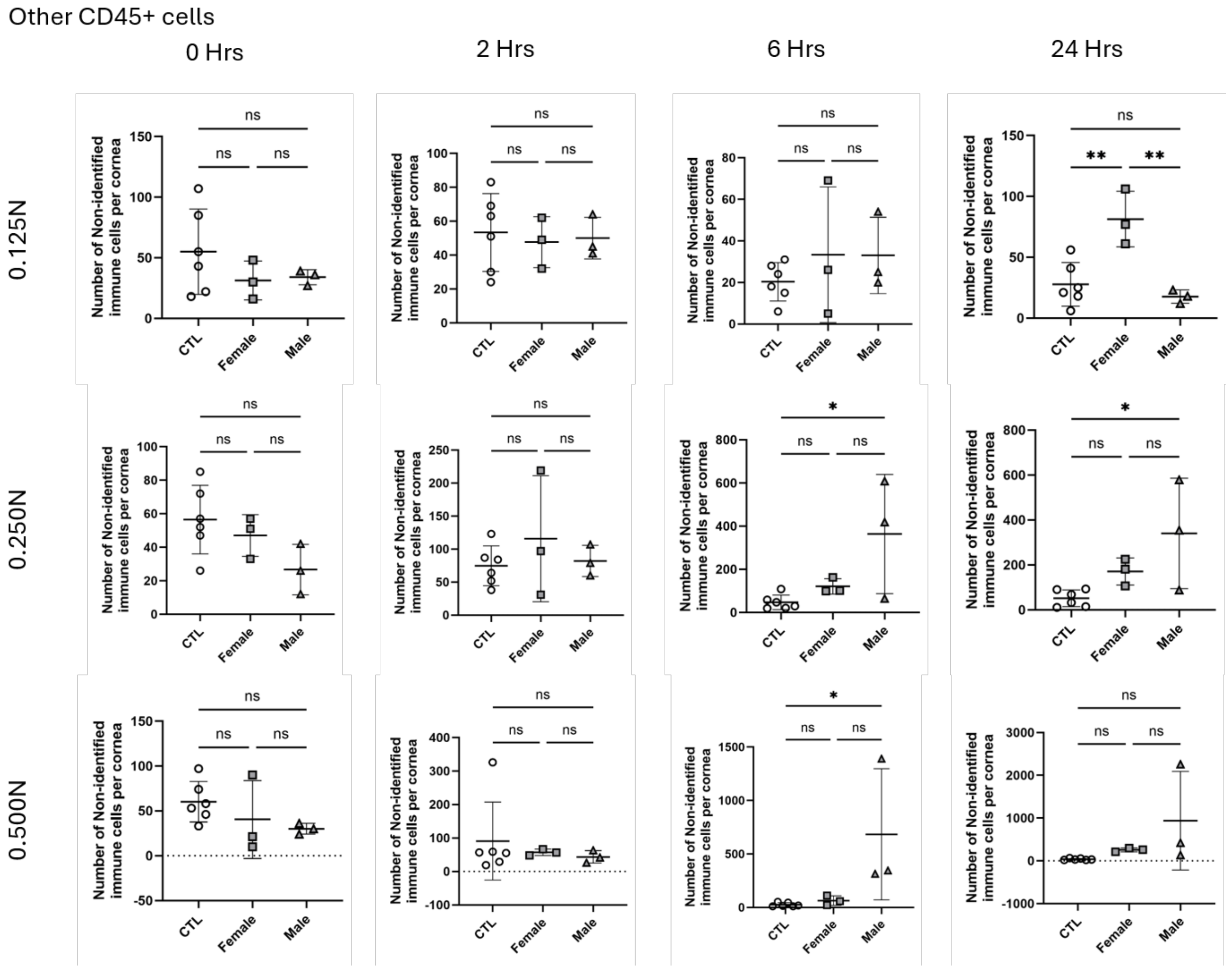


**Supplementary Figure S12**. Quantification of non-identified CD45$^+$ immune cells following corneal alkali burn. Flow cytometric analysis of residual CD45$^+$ leukocytes that were not classified within the major immune-cell populations (T cells, B cells, dendritic cells, macrophages, monocytes, neutrophils or NK cells) following corneal alkali injury. Corneas were harvested immediately (0 hours), and at 2, 6 and 24 hours after exposure to 0.125N, 0.25N or 0.5N NaOH. Data are presented separately for female and male mice and compared with untreated control (CTL) corneas. Each symbol represents one cornea; bars indicate mean ± SEM. Statistical analysis was performed using one-way ANOVA with Tukey's multiple-comparisons test. *P < 0.05, **P < 0.01; ns, not significant.

**Supplemental Table S1**. List of antibodies used for flow cytometry.

| Target | Antibody | Dilution Factor |
|---|---|---|
| CD45 | BD Pharmingen™ PerCP-Cy™5.5 Rat Anti-Mouse CD45, Clone: 30-F11, Becton, Dickinson and Company, 550994 | 1/500 |
| Ly-6C | APC/Cyanine7 anti-mouse Ly-6C Antibody, Clone: HK1.4, Biolegend, 128025 | 1/500 |
| Ly-6G | PE/Cyanine7 anti-mouse Ly-6G Antibody, Clone: 1A8, Biolegend, 127617 | 1/500 |
| CD11b | Brilliant Violet 785™ anti-mouse/human CD11b Antibody, Clone: M1/70, Biolegend, 101243 | 1/500 |
| CD11c | Brilliant Violet 650™ anti-mouse CD11c, Clone: N418, Biolegend, 117339 | 1/250 |
| F4/80 | BD Horizon™ PE-CF594 Rat Anti-Mouse F4/80, Clone: T45-2342, Becton, Dickinson and Company, 565613 | 1/250 |
| CD3e | Brilliant Violet 421™ anti-mouse CD3ε Antibody, Clone: 145-2C11, Biolegend, 100335 | 1/500 |
| CD19 | PE anti-mouse CD19 Antibody, Clone: 6D5, Biolegend, 115507 | 1/500 |
| NK1.1 | APC anti-mouse NK-1.1 Antibody, Clone: PK136, Biolegend, 108709 | 1/500 |

**Supplemental Table S2**. Statistical analysis of immune cells in the cornea.

| Figure ID | Samples being compared | Mean diff. | 95.00% CI of diff. | Below threshold? | Summary | Adjusted P Value |
|---|---|---|---|---|---|---|
| Fig. 7a, 0.125N | CTL vs. 0 Hr | 29.08 | -150.0 to 208.2 | No | ns | 0.9903 |
| | CTL vs. 2 Hrs | -127.1 | -306.2 to 52.00 | No | ns | 0.2739 |
| | CTL vs. 6 Hrs | 68.08 | -111.0 to 247.2 | No | ns | 0.8146 |
| | CTL vs. 24 Hrs | -173.3 | -352.3 to 5.838 | No | ns | 0.0622 |
| | 0 Hr vs. 2 Hrs | -156.2 | -382.7 to 70.36 | No | ns | 0.3013 |
| | 0 Hr vs. 6 Hrs | 39 | -187.5 to 265.5 | No | ns | 0.9879 |
| | 0 Hr vs. 24 Hrs | -202.3 | -428.9 to 24.20 | No | ns | 0.0999 |
| | 2 Hrs vs. 6 Hrs | 195.2 | -31.36 to 421.7 | No | ns | 0.121 |
| | 2 Hrs vs. 24 Hrs | -46.17 | -272.7 to 180.4 | No | ns | 0.9773 |
| | 6 Hrs vs. 24 Hrs | -241.3 | -467.9 to -14.80 | Yes | * | 0.0317 |
| Fig. 7a, 0.25N | CTL vs. 0 Hr | -94.25 | -590.4 to 401.9 | No | ns | 0.9825 |
| | CTL vs. 2 Hrs | -550.6 | -1047 to -54.42 | Yes | * | 0.023 |
| | CTL vs. 6 Hrs | -908.9 | -1405 to -412.8 | Yes | **** | <0.0001 |
| | CTL vs. 24 Hrs | -884.6 | -1381 to -388.4 | Yes | **** | <0.0001 |
| | 0 Hr vs. 2 Hrs | -456.3 | -1084 to 171.3 | No | ns | 0.2516 |
| | 0 Hr vs. 6 Hrs | -814.7 | -1442 to -187.1 | Yes | ** | 0.0053 |
| | 0 Hr vs. 24 Hrs | -790.3 | -1418 to -162.7 | Yes | ** | 0.0072 |

| | | | | | | |
|---|---|---|---|---|---|---|
| | 2 Hrs vs. 6 Hrs | -358.3 | -985.9 to 269.3 | No | ns | 0.4898 |
| | 2 Hrs vs. 24 Hrs | -334 | -961.6 to 293.6 | No | ns | 0.5586 |
| | 6 Hrs vs. 24 Hrs | 24.33 | -603.3 to 651.9 | No | ns | >0.9999 |
| Fig. 7a, 0.5N | CTL vs. 0 Hr | -17.25 | -2038 to 2004 | No | ns | >0.9999 |
| | CTL vs. 2 Hrs | -301.4 | -2322 to 1719 | No | ns | 0.993 |
| | CTL vs. 6 Hrs | -2236 | -4257 to -215.0 | Yes | * | 0.0235 |
| | CTL vs. 24 Hrs | -2427 | -4448 to -406.4 | Yes | * | 0.0115 |
| | 0 Hr vs. 2 Hrs | -284.2 | -2840 to 2272 | No | ns | 0.9977 |
| | 0 Hr vs. 6 Hrs | -2219 | -4775 to 337.6 | No | ns | 0.1165 |
| | 0 Hr vs. 24 Hrs | -2410 | -4966 to 146.2 | No | ns | 0.0731 |
| | 2 Hrs vs. 6 Hrs | -1935 | -4491 to 621.7 | No | ns | 0.2166 |
| | 2 Hrs vs. 24 Hrs | -2126 | -4682 to 430.4 | No | ns | 0.144 |
| | 6 Hrs vs. 24 Hrs | -191.3 | -2748 to 2365 | No | ns | 0.9995 |
| Fig. 7b, 0.125N | CTL vs. 0 Hr | 17.5 | -41.91 to 76.91 | No | ns | 0.917 |
| | CTL vs. 2 Hrs | -16.67 | -76.08 to 42.74 | No | ns | 0.9296 |
| | CTL vs. 6 Hrs | 39.83 | -19.58 to 99.24 | No | ns | 0.3283 |
| | CTL vs. 24 Hrs | -8.667 | -68.08 to 50.74 | No | ns | 0.9935 |
| | 0 Hr vs. 2 Hrs | -34.17 | -109.3 to 40.98 | No | ns | 0.696 |
| | 0 Hr vs. 6 Hrs | 22.33 | -52.81 to 97.48 | No | ns | 0.9145 |
| | 0 Hr vs. 24 Hrs | -26.17 | -101.3 to 48.98 | No | ns | 0.8578 |
| | 2 Hrs vs. 6 Hrs | 56.5 | -18.65 to 131.6 | No | ns | 0.2222 |
| | 2 Hrs vs. 24 Hrs | 8 | -67.15 to 83.15 | No | ns | 0.9981 |
| | 6 Hrs vs. 24 Hrs | -48.5 | -123.6 to 26.65 | No | ns | 0.3662 |
| Fig. 7b, 0.25N | CTL vs. 0 Hr | -47.33 | -193.4 to 98.73 | No | ns | 0.8866 |
| | CTL vs. 2 Hrs | -208 | -354.1 to -61.94 | Yes | ** | 0.0018 |
| | CTL vs. 6 Hrs | -224.7 | -370.7 to -78.61 | Yes | *** | 0.0007 |
| | CTL vs. 24 Hrs | -119.7 | -265.7 to 26.39 | No | ns | 0.1545 |
| | 0 Hr vs. 2 Hrs | -160.7 | -345.4 to 24.08 | No | ns | 0.1153 |
| | 0 Hr vs. 6 Hrs | -177.3 | -362.1 to 7.418 | No | ns | 0.0654 |
| | 0 Hr vs. 24 Hrs | -72.33 | -257.1 to 112.4 | No | ns | 0.798 |
| | 2 Hrs vs. 6 Hrs | -16.67 | -201.4 to 168.1 | No | ns | 0.999 |
| | 2 Hrs vs. 24 Hrs | 88.33 | -96.42 to 273.1 | No | ns | 0.6552 |
| | 6 Hrs vs. 24 Hrs | 105 | -79.75 to 289.8 | No | ns | 0.4944 |
| Fig. 7b, 0.5N | CTL vs. 0 Hr | 43.5 | -140.5 to 227.5 | No | ns | 0.9612 |
| | CTL vs. 2 Hrs | -168 | -352.0 to 16.03 | No | ns | 0.0884 |
| | CTL vs. 6 Hrs | -326.3 | -510.4 to -142.3 | Yes | **** | <0.0001 |
| | CTL vs. 24 Hrs | -356 | -540.0 to -172.0 | Yes | **** | <0.0001 |
| | 0 Hr vs. 2 Hrs | -211.5 | -444.3 to 21.28 | No | ns | 0.0908 |
| | 0 Hr vs. 6 Hrs | -369.8 | -602.6 to -137.1 | Yes | *** | 0.0004 |
| | 0 Hr vs. 24 Hrs | -399.5 | -632.3 to -166.7 | Yes | *** | 0.0001 |
| | 2 Hrs vs. 6 Hrs | -158.3 | -391.1 to 74.45 | No | ns | 0.3143 |
| | 2 Hrs vs. 24 Hrs | -188 | -420.8 to 44.78 | No | ns | 0.1649 |
| | 6 Hrs vs. 24 Hrs | -29.67 | -262.4 to 203.1 | No | ns | 0.9962 |
| | CTL vs. 0 Hr | -20.79 | -54.16 to 12.58 | No | ns | 0.4017 |
| | CTL vs. 2 Hrs | 4.375 | -29.00 to 37.75 | No | ns | 0.9957 |

| | | | | | | |
|---|---|---|---|---|---|---|
| Fig. 7c, 0.125N | CTL vs. 6 Hrs | 27.71 | -5.662 to 61.08 | No | ns | 0.1451 |
| | CTL vs. 24 Hrs | 26.54 | -6.829 to 59.91 | No | ns | 0.1764 |
| | 0 Hr vs. 2 Hrs | 25.17 | -17.04 to 67.38 | No | ns | 0.4463 |
| | 0 Hr vs. 6 Hrs | 48.5 | 6.289 to 90.71 | Yes | * | 0.0171 |
| | 0 Hr vs. 24 Hrs | 47.33 | 5.122 to 89.54 | Yes | * | 0.0211 |
| | 2 Hrs vs. 6 Hrs | 23.33 | -18.88 to 65.54 | No | ns | 0.5219 |
| | 2 Hrs vs. 24 Hrs | 22.17 | -20.04 to 64.38 | No | ns | 0.5712 |
| | 6 Hrs vs. 24 Hrs | -1.167 | -43.38 to 41.04 | No | ns | >0.9999 |
| Fig. 7c, 0.25N | CTL vs. 0 Hr | -23.46 | -60.79 to 13.87 | No | ns | 0.3931 |
| | CTL vs. 2 Hrs | 7.542 | -29.79 to 44.87 | No | ns | 0.978 |
| | CTL vs. 6 Hrs | 25.21 | -12.12 to 62.54 | No | ns | 0.3214 |
| | CTL vs. 24 Hrs | 26.04 | -11.29 to 63.37 | No | ns | 0.29 |
| | 0 Hr vs. 2 Hrs | 31 | -16.22 to 78.22 | No | ns | 0.3492 |
| | 0 Hr vs. 6 Hrs | 48.67 | 1.444 to 95.89 | Yes | * | 0.0405 |
| | 0 Hr vs. 24 Hrs | 49.5 | 2.277 to 96.72 | Yes | * | 0.0358 |
| | 2 Hrs vs. 6 Hrs | 17.67 | -29.56 to 64.89 | No | ns | 0.8232 |
| | 2 Hrs vs. 24 Hrs | 18.5 | -28.72 to 65.72 | No | ns | 0.7976 |
| | 6 Hrs vs. 24 Hrs | 0.8333 | -46.39 to 48.06 | No | ns | >0.9999 |
| Fig. 7c, 0.5N | CTL vs. 0 Hr | 6.208 | -26.16 to 38.58 | No | ns | 0.9819 |
| | CTL vs. 2 Hrs | 13.21 | -19.16 to 45.58 | No | ns | 0.7727 |
| | CTL vs. 6 Hrs | 27.88 | -4.493 to 60.24 | No | ns | 0.1212 |
| | CTL vs. 24 Hrs | 26.71 | -5.659 to 59.08 | No | ns | 0.1495 |
| | 0 Hr vs. 2 Hrs | 7 | -33.94 to 47.94 | No | ns | 0.9882 |
| | 0 Hr vs. 6 Hrs | 21.67 | -19.28 to 62.61 | No | ns | 0.564 |
| | 0 Hr vs. 24 Hrs | 20.5 | -20.44 to 61.44 | No | ns | 0.615 |
| | 2 Hrs vs. 6 Hrs | 14.67 | -26.28 to 55.61 | No | ns | 0.8448 |
| | 2 Hrs vs. 24 Hrs | 13.5 | -27.44 to 54.44 | No | ns | 0.8802 |
| | 6 Hrs vs. 24 Hrs | -1.167 | -42.11 to 39.78 | No | ns | >0.9999 |
| Fig. 7d, 0.125N | CTL vs. 0 Hr | 9.625 | -9.801 to 29.05 | No | ns | 0.6244 |
| | CTL vs. 2 Hrs | -7.208 | -26.63 to 12.22 | No | ns | 0.8274 |
| | CTL vs. 6 Hrs | -17.54 | -36.97 to 1.884 | No | ns | 0.094 |
| | CTL vs. 24 Hrs | -19.38 | - 38.80 to 0.05101 | No | ns | 0.0509 |
| | 0 Hr vs. 2 Hrs | -16.83 | -41.41 to 7.739 | No | ns | 0.3073 |
| | 0 Hr vs. 6 Hrs | -27.17 | -51.74 to -2.594 | Yes | * | 0.0237 |
| | 0 Hr vs. 24 Hrs | -29 | -53.57 to -4.428 | Yes | * | 0.0135 |
| | 2 Hrs vs. 6 Hrs | -10.33 | -34.91 to 14.24 | No | ns | 0.7529 |
| | 2 Hrs vs. 24 Hrs | -12.17 | -36.74 to 12.41 | No | ns | 0.625 |
| | 6 Hrs vs. 24 Hrs | -1.833 | -26.41 to 22.74 | No | ns | 0.9995 |
| Fig. 7d, 0.25N | CTL vs. 0 Hr | 4.125 | -14.70 to 22.95 | No | ns | 0.9704 |
| | CTL vs. 2 Hrs | -4.542 | -23.36 to 14.28 | No | ns | 0.9582 |
| | CTL vs. 6 Hrs | -53.54 | -72.36 to -34.72 | Yes | **** | <0.0001 |
| | CTL vs. 24 Hrs | -33.71 | -52.53 to -14.89 | Yes | **** | <0.0001 |
| | 0 Hr vs. 2 Hrs | -8.667 | -32.47 to 15.14 | No | ns | 0.837 |
| | 0 Hr vs. 6 Hrs | -57.67 | -81.47 to -33.86 | Yes | **** | <0.0001 |

| | | | | | | |
|---|---|---|---|---|---|---|
| | 0 Hr vs. 24 Hrs | -37.83 | -61.64 to -14.03 | Yes | *** | 0.0004 |
| | 2 Hrs vs. 6 Hrs | -49 | -72.81 to -25.19 | Yes | **** | <0.0001 |
| | 2 Hrs vs. 24 Hrs | -29.17 | -52.97 to -5.361 | Yes | ** | 0.0095 |
| | 6 Hrs vs. 24 Hrs | 19.83 | -3.972 to 43.64 | No | ns | 0.1428 |
| Fig. 7d, 0.5N | CTL vs. 0 Hr | 6.625 | -24.22 to 37.47 | No | ns | 0.9725 |
| | CTL vs. 2 Hrs | -4.208 | -35.06 to 26.64 | No | ns | 0.995 |
| | CTL vs. 6 Hrs | -76.21 | -107.1 to -45.36 | Yes | **** | <0.0001 |
| | CTL vs. 24 Hrs | -52.71 | -83.56 to -21.86 | Yes | *** | 0.0001 |
| | 0 Hr vs. 2 Hrs | -10.83 | -49.85 to 28.19 | No | ns | 0.932 |
| | 0 Hr vs. 6 Hrs | -82.83 | -121.9 to -43.81 | Yes | **** | <0.0001 |
| | 0 Hr vs. 24 Hrs | -59.33 | -98.35 to -20.31 | Yes | *** | 0.0008 |
| | 2 Hrs vs. 6 Hrs | -72 | -111.0 to -32.98 | Yes | **** | <0.0001 |
| | 2 Hrs vs. 24 Hrs | -48.5 | -87.52 to -9.480 | Yes | ** | 0.0083 |
| | 6 Hrs vs. 24 Hrs | 23.5 | -15.52 to 62.52 | No | ns | 0.4361 |
| Fig. 7e, 0.125N | CTL vs. 0 Hr | 10.13 | -15.14 to 35.39 | No | ns | 0.7839 |
| | CTL vs. 2 Hrs | 0.2917 | -24.97 to 25.55 | No | ns | >0.9999 |
| | CTL vs. 6 Hrs | 0.2917 | -24.97 to 25.55 | No | ns | >0.9999 |
| | CTL vs. 24 Hrs | 3.292 | -21.97 to 28.55 | No | ns | 0.9958 |
| | 0 Hr vs. 2 Hrs | -9.833 | -41.79 to 22.12 | No | ns | 0.9041 |
| | 0 Hr vs. 6 Hrs | -9.833 | -41.79 to 22.12 | No | ns | 0.9041 |
| | 0 Hr vs. 24 Hrs | -6.833 | -38.79 to 25.12 | No | ns | 0.9729 |
| | 2 Hrs vs. 6 Hrs | 0 | -31.95 to 31.95 | No | ns | >0.9999 |
| | 2 Hrs vs. 24 Hrs | 3 | -28.95 to 34.95 | No | ns | 0.9988 |
| | 6 Hrs vs. 24 Hrs | 3 | -28.95 to 34.95 | No | ns | 0.9988 |
| Fig. 7e, 0.25N | CTL vs. 0 Hr | 25.96 | -0.7490 to 52.67 | No | ns | 0.0604 |
| | CTL vs. 2 Hrs | 3.792 | -22.92 to 30.50 | No | ns | 0.9942 |
| | CTL vs. 6 Hrs | 8.125 | -18.58 to 34.83 | No | ns | 0.9077 |
| | CTL vs. 24 Hrs | 6.125 | -20.58 to 32.83 | No | ns | 0.9652 |
| | 0 Hr vs. 2 Hrs | -22.17 | -55.95 to 11.62 | No | ns | 0.3496 |
| | 0 Hr vs. 6 Hrs | -17.83 | -51.62 to 15.95 | No | ns | 0.5663 |
| | 0 Hr vs. 24 Hrs | -19.83 | -53.62 to 13.95 | No | ns | 0.4619 |
| | 2 Hrs vs. 6 Hrs | 4.333 | -29.45 to 38.12 | No | ns | 0.9961 |
| | 2 Hrs vs. 24 Hrs | 2.333 | -31.45 to 36.12 | No | ns | 0.9997 |
| | 6 Hrs vs. 24 Hrs | -2 | -35.78 to 31.78 | No | ns | 0.9998 |
| Fig. 7e, 0.5N | CTL vs. 0 Hr | 27.46 | 6.074 to 48.84 | Yes | ** | 0.0059 |
| | CTL vs. 2 Hrs | 26.79 | 5.407 to 48.18 | Yes | ** | 0.0076 |
| | CTL vs. 6 Hrs | 28.46 | 7.074 to 49.84 | Yes | ** | 0.0041 |
| | CTL vs. 24 Hrs | 18.79 | -2.593 to 40.18 | No | ns | 0.1091 |
| | 0 Hr vs. 2 Hrs | -0.6667 | -27.72 to 26.38 | No | ns | >0.9999 |
| | 0 Hr vs. 6 Hrs | 1 | -26.05 to 28.05 | No | ns | >0.9999 |
| | 0 Hr vs. 24 Hrs | -8.667 | -35.72 to 18.38 | No | ns | 0.8907 |
| | 2 Hrs vs. 6 Hrs | 1.667 | -25.38 to 28.72 | No | ns | 0.9998 |
| | 2 Hrs vs. 24 Hrs | -8 | -35.05 to 19.05 | No | ns | 0.9159 |
| | 6 Hrs vs. 24 Hrs | -9.667 | -36.72 to 17.38 | No | ns | 0.846 |

| | | | | | | |
|---|---|---|---|---|---|---|
| Fig. 7f, 0.125N | CTL vs. 0 Hr | 3.667 | -32.51 to 39.84 | No | ns | 0.9984 |
| | CTL vs. 2 Hrs | -27 | -63.18 to 9.176 | No | ns | 0.2285 |
| | CTL vs. 6 Hrs | 1.667 | -34.51 to 37.84 | No | ns | >0.9999 |
| | CTL vs. 24 Hrs | -37.17 | -73.34 to -0.9910 | Yes | * | 0.0414 |
| | 0 Hr vs. 2 Hrs | -30.67 | -76.43 to 15.09 | No | ns | 0.3287 |
| | 0 Hr vs. 6 Hrs | -2 | -47.76 to 43.76 | No | ns | >0.9999 |
| | 0 Hr vs. 24 Hrs | -40.83 | -86.59 to 4.926 | No | ns | 0.1004 |
| | 2 Hrs vs. 6 Hrs | 28.67 | -17.09 to 74.43 | No | ns | 0.3961 |
| | 2 Hrs vs. 24 Hrs | -10.17 | -55.93 to 35.59 | No | ns | 0.9689 |
| | 6 Hrs vs. 24 Hrs | -38.83 | -84.59 to 6.926 | No | ns | 0.1305 |
| Fig. 7f, 0.25N | CTL vs. 0 Hr | 3.667 | -179.8 to 187.2 | No | ns | >0.9999 |
| | CTL vs. 2 Hrs | -203 | -386.5 to -19.50 | Yes | * | 0.0236 |
| | CTL vs. 6 Hrs | -162.5 | -346.0 to 21.00 | No | ns | 0.1047 |
| | CTL vs. 24 Hrs | -278.5 | -462.0 to -95.00 | Yes | *** | 0.0008 |
| | 0 Hr vs. 2 Hrs | -206.7 | -438.8 to 25.45 | No | ns | 0.1017 |
| | 0 Hr vs. 6 Hrs | -166.2 | -398.3 to 65.95 | No | ns | 0.2658 |
| | 0 Hr vs. 24 Hrs | -282.2 | -514.3 to -50.05 | Yes | * | 0.0102 |
| | 2 Hrs vs. 6 Hrs | 40.5 | -191.6 to 272.6 | No | ns | 0.9873 |
| | 2 Hrs vs. 24 Hrs | -75.5 | -307.6 to 156.6 | No | ns | 0.8853 |
| | 6 Hrs vs. 24 Hrs | -116 | -348.1 to 116.1 | No | ns | 0.6167 |
| Fig. 7f, 0.5N | CTL vs. 0 Hr | -89.5 | -400.1 to 221.1 | No | ns | 0.9229 |
| | CTL vs. 2 Hrs | -64.17 | -374.8 to 246.5 | No | ns | 0.9762 |
| | CTL vs. 6 Hrs | -113.5 | -424.1 to 197.1 | No | ns | 0.8352 |
| | CTL vs. 24 Hrs | -702.8 | -1013 to -392.2 | Yes | **** | <0.0001 |
| | 0 Hr vs. 2 Hrs | 25.33 | -367.6 to 418.2 | No | ns | 0.9997 |
| | 0 Hr vs. 6 Hrs | -24 | -416.9 to 368.9 | No | ns | 0.9998 |
| | 0 Hr vs. 24 Hrs | -613.3 | -1006 to -220.4 | Yes | *** | 0.0006 |
| | 2 Hrs vs. 6 Hrs | -49.33 | -442.2 to 343.6 | No | ns | 0.9964 |
| | 2 Hrs vs. 24 Hrs | -638.7 | -1032 to -245.8 | Yes | *** | 0.0003 |
| | 6 Hrs vs. 24 Hrs | -589.3 | -982.2 to -196.4 | Yes | *** | 0.001 |
| Fig. 8a, 0 hours | CTL vs. Female | -81.83 | -446.1 to 282.5 | No | ns | 0.8093 |
| | CTL vs. Male | -52.5 | -416.8 to 311.8 | No | ns | 0.9154 |
| | Female vs. Male | 29.33 | -391.3 to 450.0 | No | ns | 0.9794 |
| Fig. 8a, 2 hours | CTL vs. Female | -549.5 | -1271 to 171.8 | No | ns | 0.1391 |
| | CTL vs. Male | -356.2 | -1078 to 365.2 | No | ns | 0.3912 |
| | Female vs. Male | 193.3 | -639.6 to 1026 | No | ns | 0.7981 |
| Fig. 8b, 0 hours | CTL vs. Female | 0.1667 | -1.999 to 2.333 | No | ns | 0.9749 |
| | CTL vs. Male | 0.1667 | -1.999 to 2.333 | No | ns | 0.9749 |
| | Female vs. Male | 0 | -2.501 to 2.501 | No | ns | >0.9999 |
| Fig. 8b, 2 hours | CTL vs. Female | -358.8 | -772.4 to 54.71 | No | ns | 0.0887 |
| | CTL vs. Male | -40.17 | -453.7 to 373.4 | No | ns | 0.9604 |
| | Female vs. Male | 318.7 | -158.9 to 796.2 | No | ns | 0.2046 |
| | CTL vs. Female | -10.67 | -146.2 to 124.8 | No | ns | 0.9738 |
| | CTL vs. Male | -90.33 | -225.8 to 45.17 | No | ns | 0.2051 |

| | | | | | | |
|---|---|---|---|---|---|---|
| Fig. 8c, 0 hours | Female vs. Male | -79.67 | -236.1 to 76.80 | No | ns | 0.3709 |
| Fig. 8c, 2 hours | CTL vs. Female | -18.17 | -81.35 to 45.01 | No | ns | 0.7107 |
| | CTL vs. Male | -79.17 | -142.3 to -15.99 | Yes | * | 0.0167 |
| | Female vs. Male | -61 | -134.0 to 11.95 | No | ns | 0.1015 |
| Fig. 8d, 0 hours | CTL vs. Female | 14.83 | -16.09 to 45.75 | No | ns | 0.4102 |
| | CTL vs. Male | 28.5 | -2.419 to 59.42 | No | ns | 0.0703 |
| | Female vs. Male | 13.67 | -22.04 to 49.37 | No | ns | 0.5554 |
| Fig. 8d, 2 hours | CTL vs. Female | 8.167 | -34.92 to 51.26 | No | ns | 0.8593 |
| | CTL vs. Male | 47.5 | 4.409 to 90.59 | Yes | * | 0.032 |
| | Female vs. Male | 39.33 | -10.42 to 89.09 | No | ns | 0.1233 |
| Fig. 8e, 0 hours | CTL vs. Female | 3.167 | -18.30 to 24.63 | No | ns | 0.9116 |
| | CTL vs. Male | 8.5 | -12.97 to 29.97 | No | ns | 0.5346 |
| | Female vs. Male | 5.333 | -19.46 to 30.12 | No | ns | 0.8232 |
| Fig. 8e, 2 hours | CTL vs. Female | 3.333 | -15.81 to 22.47 | No | ns | 0.8795 |
| | CTL vs. Male | 4 | -15.14 to 23.14 | No | ns | 0.8322 |
| | Female vs. Male | 0.6667 | -21.44 to 22.77 | No | ns | 0.9961 |
| Fig. 8f, 0 hours | CTL vs. Female | -59.33 | -214.5 to 95.81 | No | ns | 0.556 |
| | CTL vs. Male | -72.33 | -227.5 to 82.81 | No | ns | 0.4291 |
| | Female vs. Male | -13 | -192.1 to 166.1 | No | ns | 0.9777 |
| Fig. 8f, 2 hours | CTL vs. Female | -140.2 | -426.9 to 146.5 | No | ns | 0.3977 |
| | CTL vs. Male | -261.2 | -547.9 to 25.55 | No | ns | 0.0736 |
| | Female vs. Male | -121 | -452.1 to 210.1 | No | ns | 0.5833 |
| Fig. 8g, 0 hours | CTL vs. Female | -39.33 | -129.8 to 51.18 | No | ns | 0.4752 |
| | CTL vs. Male | 42 | -48.51 to 132.5 | No | ns | 0.4322 |
| | Female vs. Male | 81.33 | -23.18 to 185.8 | No | ns | 0.1298 |
| Fig. 8g, 2 hours | CTL vs. Female | -1.667 | -24.75 to 21.42 | No | ns | 0.9779 |
| | CTL vs. Male | -3.667 | -26.75 to 19.42 | No | ns | 0.8985 |
| | Female vs. Male | -2 | -28.66 to 24.66 | No | ns | 0.9762 |
| Fig. 8h, 0 hours | CTL vs. Female | -0.1667 | -1.969 to 1.636 | No | ns | 0.9641 |
| | CTL vs. Male | 0.5 | -1.302 to 2.302 | No | ns | 0.727 |
| | Female vs. Male | 0.6667 | -1.414 to 2.748 | No | ns | 0.657 |
| Fig. 8h, 2 hours | CTL vs. Female | -1.333 | -19.74 to 17.08 | No | ns | 0.9778 |
| | CTL vs. Male | -16 | -34.41 to 2.411 | No | ns | 0.0882 |
| | Female vs. Male | -14.67 | -35.93 to 6.592 | No | ns | 0 |
